# Mitogen-independent cell cycle progression in B lymphocytes

**DOI:** 10.1101/796904

**Authors:** Amit Singh, Matthew H. Spitzer, Jaimy P. Joy, Mary Kaileh, Xiang Qiu, Garry P. Nolan, Ranjan Sen

## Abstract

The canonical view of the cell cycle posits that G1 progression signals are essential after each mitosis to enter S phase. A subset of tumor cells bypass this requirement and progress to the next cell division in the absence of continued signaling. B and T lymphocytes of the adaptive immune system undergo a proliferative burst, termed clonal expansion, to generate pools of antigen specific cells for effective immunity. There is evidence that rules for lymphocyte cell division digress from the canonical model. Here we show that B lymphocytes sustain several rounds of mitogen-independent cell division following the first mitosis. Such division is driven by unique characteristics of the post mitotic G1 phase and limited by extensive cell death that can be circumvented by appropriate anti-apoptotic signals. An essential component for continued cell division is Birc5 (survivin), a protein associated with chromosome segregation in G2/M. Our observation provides direct evidence for Pardee’s hypothesis that retention of features of G2M in post-mitotic cells could trigger further cell cycle progression. The partially active G1 phase and propensity for apoptosis that is inherited after each division may permit rapid burst of proliferation and cell death that are hallmarks of immune responses.

## Introduction

Activation of antigen receptors of B and T lymphocytes is one key signal that initiates responses to specific pathogens^1, 2^. Antigen recognition selects a tiny proportion of lymphocytes which are induced to proliferate and execute an immune response. This proliferative phase, termed as clonal expansion, determines effectiveness of adaptive immune responses. Additionally, antigen non-specific co-stimulatory signals provided by innate immune cells contribute to the magnitude of clonal expansion^3, 4^. Regulation of lymphocyte proliferation has been studied by treating cells *ex vivo* with one or more signals followed by analyses of proliferative responses. In recent years, elegant experimental schemes have been coupled with rigorous computational studies to gain deeper insight into these processes^5–7^. Two related questions lie at the heart of these studies. First, what determines how many lymphocytes out of the pre-existing pool will proliferate? Second, how do lymphocytes count the number of divisions so as to have an effective, but limited, proliferative phase?

The prevailing view is that lymphocytes are induced to proliferate when the cumulative response to in coming signals crosses an activation threshold. The proto-oncogene Myc features prominently during proliferative phases of immune responses *in vivo*. For example, Myc is required for generation and maintenance of germinal centers (GC) within which clonally expanded B cells undergo changes essential for effective immunity and generation of memory^8, 9^. The close relationship of Myc to B cell proliferation is further highlighted by its well-established role in B cell lymphomagenesis^10, 11^. Additionally, combined absence of c-Myc and n-Myc abrogates proliferation of B lymphocytes *in vitro*^12^. For lymphocytes, the concept of division destiny (DD) has been invoked to signify the number of times a cell divides in response to mitogenic signals^13, 14^. DD is related to the mean division number (MDN), which is the average number of cell divisions made over a defined time period. It has been shown that DD is proportionate to levels of Myc that are induced in response to the initiating signal(s)^15^. Similarly, the extent of cell division in GCs has been recently shown to be proportional to Myc levels^16^.

The majority of studies of cell division in lymphocytes are carried out with persistent mitogenic stimulation, presumably reflecting our bias that mitogenic re-stimulation is necessary to get multiple rounds of cell division for robust clonal expansion. However, it remains unclear whether clonal expansion *in vivo* requires persistent antigenic stimulation. There is good evidence that signaling and proliferative phases can be spatially segregated. This is most obvious in the GC, where B cell activation via T cell help has been shown to be restricted to the GC light zone (LZ), whereas bulk of B cell proliferation occurs in the GC dark zone (DZ)^17, 18^. Recent studies attribute this to synergistic effect of BCR and CD40 signals in the LZ for optimal Myc induction^19^. The simplest interpretation of this distinction is that GC B cells can undergo multiple rounds of division without requiring continued mitogenic signaling. There is also tantalizing evidence that naive B cells can undergo such signal-independent division *ex vivo*, which suggests that this form of division may also be involved in clonal expansion^15^. Additionally, CD8^+^ T cells have been shown to undergo several rounds of division in the absence of persistent signaling^20, 21^, suggesting that this may be a common feature of cell cycle regulation in lymphocytes. Despite its uniqueness and likely physiological relevance, this form of proliferation has received scant attention.

The decision-making process after mitosis that shunts daughter cells into a G0 state or permits their re-entry into G1 has elicited much interest. The classical view, obtained largely from population analyses of fibroblasts, posits that a mitogenic signal is required at the end of each mitosis in order for cells to progress through the next G1 phase. However, analyses of tumor cell lines and, more recently, untransformed mouse embryonic stem cells (ES), have shown that a subset of cells within a population retains the ability to undergo G1 progression without additional signaling^22, 23^. Molecules that regulate this form of cell cycle progression include CDK2 and p53. CDK2 activity is normally down-regulated after mitosis. However, cells that maintain CDK2 activity have been shown to traverse G1 in the absence of additional mitogenic signals^22^. Similarly, down-regulation of p53 in post-mitotic G1 phase is required to permit re-entry into the cell cycle^24^. Although these players have not been extensively investigated during proliferative responses of primary lymphocytes, there is good evidence that p53 regulates proliferation of murine T lymphocytes^25^.

Here we characterize mitogen-independent proliferation in primary murine B lymphocytes. We demonstrate that, regardless of the initiating stimulus, commitment to DNA replication (S phase) programs B cells to undergo several rounds of cell division in the absence of overt mitogenic signaling. The extent of division is limited by cell death rather than by return to quiescence, and circumventing cell death permits up to 5 rounds of division in 72h. Mitogen-independent cell cycle progression is driven by unique characteristics of the G1 phase of cells that have divided once. The G1 of these cells have features of S and G2/M phases, including large cell size, low levels of p27 and phosphorylated Rb. In contrast to studies in cell lines, however, B cell division under these conditions requires CDK4/6 activity to traverse the second G1. Transcriptional and protein analyses revealed up-regulation of survivin (Birc5) in the G1 phase of B cells past the first mitosis. Inhibition of survivin function with pharmacologic inhibitors blocked G1 progression of cells undergoing mitogen-independent proliferation, but not of naïve B cells stimulated with mitogens. These observations indicate that, in contrast to textbook models of the cell cycle, B cells inherit a partially active G1 phase after cell division that permits them to move quickly to the next S phase in the absence of exogenous G1 progression signals. Our studies provide direct evidence for Pardee’s hypothesis^26^ that retention of features of G2/M in post-mitotic cells could trigger a second round of cell cycle progression. We propose that these mechanisms may assist in rapid cell division without differentiation that is required for clonal expansion in response to antigen and in B cell proliferation in the GC dark zone.

## Results

### B cells undergo mitogen-independent division

DeFranco and Paul first demonstrated that B cell mitogenic signals could be removed at a defined time before initiation of DNA synthesis without affecting the ability of cells to enter S phase^27^. In modern parlance this observation reflects the need for continuous signaling for cells to progress past the restriction point in G1^28^. The restriction point is characterized by phosphorylation of Rb, release of E2F factors, degradation of cell cycle inhibitors (p21 or p27) and inactivation of Cdh1^29^, continued signaling is not required to proceed into S phase and through mitosis. Thereafter, mitogenic signals are again required to induce G1 progression past the restriction point to achieve the second division. In contrast to this model, robust CD8^+^ T cell proliferation through several rounds of cell division has been observed even if mitogens are removed prior to the first cell division^20, 21^. Similar properties have also been attributed to B cells^30^. However, mechanisms that permit lymphocytes to deviate from the classical model have not been explored. To better understand this phenomenon, we started our analysis by characterizing the requirement for mitogenic signals for B cell proliferation.

We activated naive splenic B lymphocytes with a variety of mitogenic signals for 24h, and thereafter followed cell division in the absence of overt signaling. We found that none of these treatments induced cell division within 24h. Yet, cells progressed through multiple rounds of cell division in the next 48h regardless of whether mitogen was present continuously or not (Figure 1A). However, cells that lacked continuous mitogenic signals were considerably less viable (Figure 1A, numbers in italics). The proportion of cells that divided in the absence of mitogens differed depending on the stimulus. As previously noted^13^, persistent CpG treatment led to reduced cell size with increased division (Figure 1B). For all other stimuli, however, cell size increased with additional division even without continued stimulation (Figure 1B, green labels). We conclude that splenic B lymphocytes are programmed to divide multiple times in the absence of continued stimulation. This decision is made during the first cell cycle and is limited by cell death.

**Figure 1.**
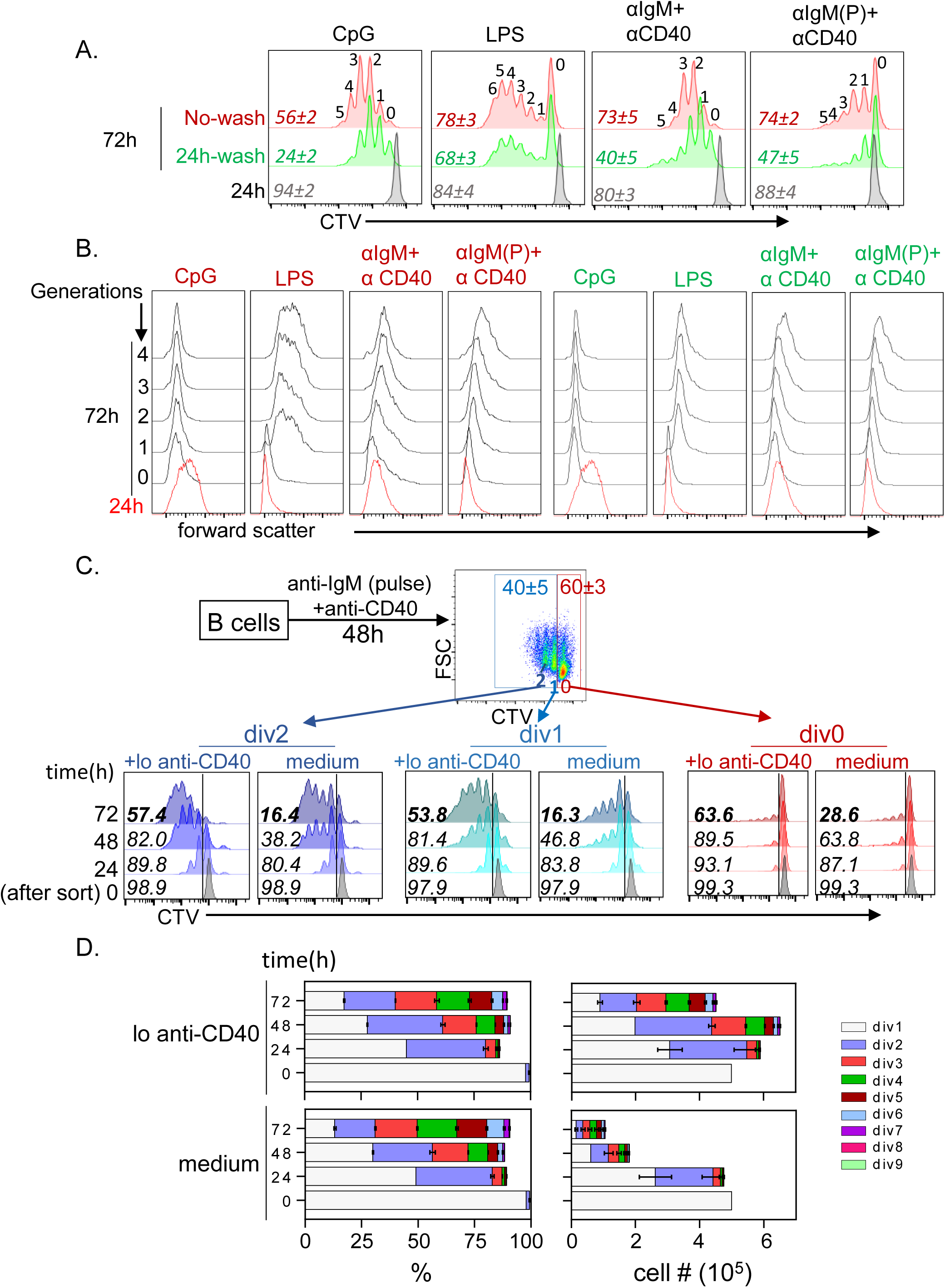
Signal-independent proliferation of murine spleen B cells. Splenic B cells were activated with CpG (50 ng/ml), LPS (5 µg/ml), F(ab’)2 anti-IgM (10 µg/ml) + anti-CD40 antibody (5 µg/ml) or pulse BCR stimulation followed by anti-CD40 treatment^29^ as indicated. After 24h half the cells were washed free of the stimulus and further incubated for an additional 48h. Remaining cells were continuously stimulated for the entire 72h period, followed by flow cytometric analysis. A) Cell division was assessed by CTV dilution at 24h (grey) and at 72h (green and red); ‘No wash’ (red) refers to cells that were continuously stimulated. Cells that were washed and incubated further in the absence of mitogen are shown in green. Numbers in italics refer to the percentage of viable cells in each culture; numbers above CTV peaks refers to the cell generations. B) Cell size was estimated by forward scatter for each cell generation at 24 and 72h. Red and green labeling refer to cells that had been continuously stimulated or washed free of stimulus at 24h, respectively. C) After 48h pulse anti-IgM plus antiCD40 activation B cells were sorted into populations of cells that had not non-divided (div0), divided once (div1) and divided twice (div2). Each population was further cultured in medium alone or in medium with 0.125 µg/ul anti-CD40 (lo anti-CD40). Cell division was accessed by CTV dilution after various times of culture indicated on the left. Numbers within histogram boxes refer to percentage of viable cells under each condition. D) Cell percentages (left) and cell numbers (right) generated after culture of div1 cells for 48h in the absence of stimulation. Cell division was determined by CTV dilution as in part C (colored bars) using proliferation modeling tool in FlowJo. Cells shown in the upper panel were treated with low anti-CD40 (lo anti-CD40) and cells shown in the lower panel were incubated in medium alone (medium). Data shown is average of three independent experiments with error bars showing SEM.

These observations raised two immediate questions. First, what determines the proportion of cells that divide without further stimulation, and second, can mitogen-independent proliferation be uncoupled from extensive cell death. To address these questions, we activated cells with a pulse of BCR crosslinking followed by anti-CD40 treatment to mimic T-dependent proliferation of antigen activated B cells ^31^. This regimen led to similar numbers of divided (labeled div1 and div2) and undivided (div0) cells after 48h, permitting further analyses of both populations (Figure 1C, top). Sorted div1, div2 or div0 cells were cultured for an additional 72h in the absence of mitogenic signals. We found that very few div0 cells divided further during culture, whereas div1 and div2 cells divided robustly despite undergoing extensive cell death (Figure 1C, labeled “medium”). Similar results were observed with other mitogens (Supplementary Figure 1A). We conclude that B cells that have completed one mitotic division gain the ability to undergo several additional rounds of mitogen-independent proliferation.

To circumvent cell death, we repeated these experiments by culturing sorted cells in BAFF-containing medium. Though treatment with BAFF averted cell death, it also drastically reduced cell division especially at later cell division stages (Supplementary Figure 1B). Taking a cue from the known properties of CD40 as a poor mitogen but an excellent inducer of cell viability^32^, we included varying concentrations of two different clones of anti-CD40 antibody during post-sort culture (Supplementary Figure 1B). We found that very low concentrations of anti-CD40 (0.125 µg/ml) substantially enhanced cell viability during mitogen-independent proliferation (Figure 1C, labeled “+lo anti-CD40”), while maintaining proliferation (Supplementary Figure 1B) and cell size (Supplementary Figure 1C). Importantly, this dose of anti-CD40 did not induce proliferation of naïve B cells (data not shown) nor of activated cells that had not divided during the first 48h (Figure 1C “div0 +lo anti-CD40”). Furthermore, low dose anti-CD40 did not change the proportions of cells that divided during post-sort culture but instead increased cell numbers after longer culture periods (Figure 1D). We conclude that low levels of CD40 provide survival signals without affecting mitogen-independent cell division, suggesting that this form of lymphocyte proliferation can generate substantial cell numbers in the presence of appropriate anti-apoptotic activity.

### Cell cycle characteristics beyond first mitosis

To uncover the basis for mitogen-independent proliferation of div1 and div2 (div12) cells, we first ruled out the possibility that div0 cells were refractory to further division. For this we treated sorted div0 cells with anti-IgM and monitored proliferation by CFSE dilution. This evoked comparable mitogen-induced cell division of both div0 and div12 cells (Supplementary Figure 2A). Thus, div0 cells are capable of division. To identify differences between div0 and div12 cells, we assayed biochemical features of different stages of the cell cycle. We found that p27 levels were lower in div12 cells (Figure 2A) whereas hyper-phosphorylated Rb (S807/S811), cMyc and phospho-CDK2 (Thr160) levels were higher in div12 compared to div0 cells (Figure 2B). Because cells in G1 constituted the majority of cells in both div0 and div12 populations (Figure 2C), we tentatively concluded that these biochemical changes reflected differences in G1 states of the respective populations. By gating directly on G1 populations we found that phosphorylated Rb (Ser807/811), phospho-CDK2 (Thr160) and c-Myc levels were significantly higher in the G1 phase of div12 cells compared to the G1 of div0 cells (Figure 2D). Div0 and div12 cells in G1 also differed in cell size and granularity (Supplementary Figure 2B and 2C). These observations demonstrate that the G1 phase of post-mitotic B cells is distinct from that of activated but non-divided B cells.

**Figure 2:**
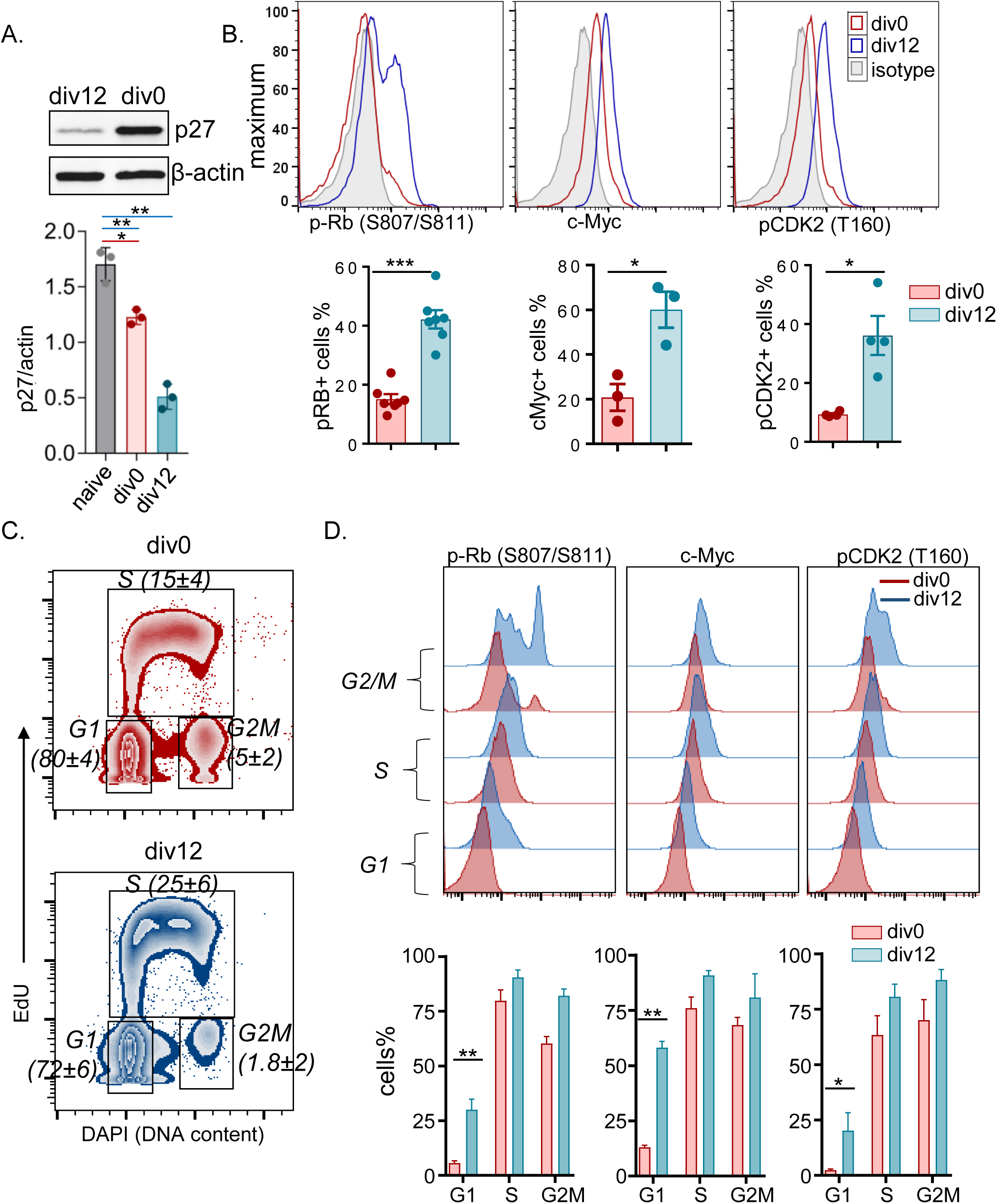
Post-mitotic B cells are enriched for markers of G1 progression. Pulse anti-IgM+anti-CD40 activated cells (48h) were purified based on CTV levels into undivided (div0) and divided (div12) populations. Cell cycle features were analyzed biochemically or by flow cytometry. A) Immunoblotting analysis of p27 levels (top). Whole cell extracts from div0 or div12 cells as indicated were fractionated by SDS-PAGE, transferred to nitrocellulose membranes which were probed with anti-p27 antibody or anti-β actin antibody for normalization. Protein levels were quantitated by densitometry and results from 3 independent experiments are represented in the bottom panel. Error bars show SEM from 3 independent experiments. B) Representative histograms showing intracellular levels of phospho-Rb (pS807/S811), c-Myc and phospho-CDK2 (pT160) in div0 (red) and div12 (blue) cells detected by flow cytometry. Percentage of cells that express indicated target proteins from several independent experiments is shown in bottom panel. Each experiment is represented by a dot and error bars show SEM. C) After 48h activation cells were pulse-labeled with EdU for 30 minutes followed by Clickit reaction for AlexaFlour labeling. Cell cycle stages in div0 and div12 cells were identified as G1 phase (2n DAPI EdU^-^), S phase (EdU^+^) and G2/M phase (4n DAPI EdU^-^). Representative zebra plots of div0 (top) and div12 (bottom) cells are shown. Numbers in parentheses indicate the percentage ±SEM of cells in each cell cycle stage derived from 6 independent experiments. D) Representative histograms of cell cycle stage-specific intracellular levels of phospho-Rb (pS807/S811), c-Myc and phospho-CDK2 (pT160) in div0 (red) and div12 (blue) cells (top). Mean protein expression from 2 to 4 independent experiments is shown in the lower panel; error bars represent SEM.

Persistent activity of CDK2 in G1 has been shown to permit a subset of tumor cells to circumvent the requirement for mitogen-induced activation of CDK4/6 for G1 progression^22^. To determine whether mitogen-independent B cell proliferation was similarly regulated, we assessed the requirement for CDK1/2 (specific for S and G2/M) and CDK4/6 (specific for G1) using well established pharmacologic inhibitors. Sorted div12 cells were cultured in the presence of either a CDK4/6 inhibitor (Palbociclib, iCdk4/6) or a CDK1/2 inhibitor (CDK1/2 Inhibitor III, iCdk1/2) for 48h, followed by analyses of cell division and cell cycle profiles. At least 3 additional cell divisions were evident in cells cultured without inhibitors (Figure 3A and quantified in 3B, labeled DMSO). By contrast, a fraction of cells treated with iCdk4/6 divided once and then accumulated in G1, whereas CDK1/2 inhibition resulted in cells piling up in S/G2 without division (Figure 3A-C). Similar patterns were observed when signal dependent proliferation was assessed after activation of div0 population with anti-IgM (Figure 3D and 3E). We inferred that pre-existing S/G2M phase cells in the div12 population, divided once in the presence of iCdk4/6; however, absence of progression to the next S phase demonstrated that mitogen-independent G1 progression required CDK4/6 activity. By contrast, iCdk1/2 prevented mitosis of pre-existing S/G2M cells within div12, resulting in a CFSE pattern similar to that of div12 cells immediately after sorting (Figure 3A). We conclude that primary B cells undergoing signal independent proliferation require CDK4/6 activity to progress through G1.

**Figure 3:**
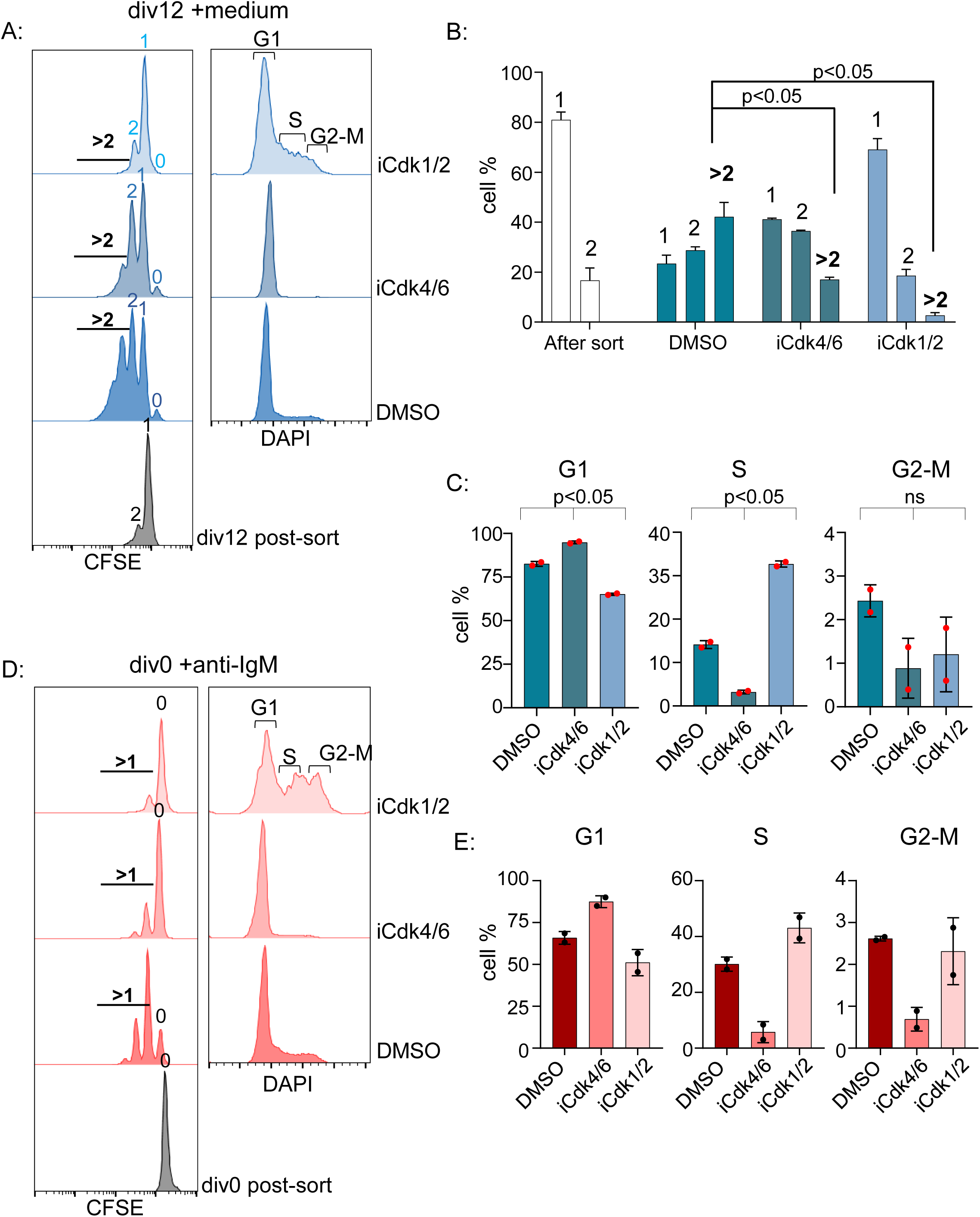
Effect of CDK inhibition on mitogen-independent B cell proliferation. Pulse anti-IgM+anti-CD40 activated cells (48h) were purified by into undivided (div0) and divided (div12) based on CFSE dilution. After separation, div12 were cultured in medium alone in the presence or absence of CDK inhibitors as indicated. Div0 cells were treated with anti-IgM in the presence or absence of the same CDK inhibitors. CDK 4/6 inhibitor (Palbociclib, iCdk4/6) was used at 1µM final concentration and CDK1/2 inhibitor (Cdk1/2 Inhibitor III, iCdk1/2) at 10µM final concentration. Proliferation was analyzed by CFSE dilution and cell cycle states by DNA content analysis using DAPI. A) Representative histograms of cell division (left) and cell cycle stages (right) measured after 48h culture of purified div12 cells in medium, with no inhibitor (labeled DMSO) or CDK inhibitors as indicated. Cell division numbers are indicated within CFSE profiles. B, C) Two independent inhibitor experiments from part A were quantified to identify percentage of cells present in each division number (B) or in each stage of the cell cycle (C). Error bars represent SEM (unpaired t-test). D) Representative histograms of cell division (left) and cell cycle stages (right) measured after 48h culture of purified div0 cells re-stimulated with anti-IgM in the presence of no inhibitor (labelled DMSO) or CDK inhibitors as indicated. Cell division numbers are indicated within CFSE profiles. E) Two independent inhibitor experiments from part D were quantified to identify percentage of cells present in each stage of the cell cycle. Error bars represent SEM (unpaired t-test).

### Differences in pre- and post-mitotic G1

We used mass cytometry by time-of-flight (CyTOF) to compare G1 states of div0 and div12 cells after 48h of mitogen-dependent culture (Figure 4A, Phase-I). Purified div0 and div12 cells were sequentially stained with iodo-deoxyuridine (IdU) and cisplatin to label S phase cells and dead cells, respectively. Following fixation, cells were labeled with antibodies directed against cell surface markers, and following permeabilization, with antibodies against cell cycle markers and signaling-related phosphoproteins as previously described^33^. Remaining div0 and div12 cells were cultured for additional 48h in medium before being similarly processed for CyTOF analysis (Figure 4A, Phase-II). A total of 28 parameters (Supplementary Figure 4A) were used to identify 200 clusters from the 3 conditions from Phase-I using SPADE algorithm. Each cell population assayed in Phase-I was distinct and relatively homogenous (Figure 4B). To probe cell cycle status of each population we used phosphorylated histone H3 (H3 Ser10) to identify M-phase cells, cyclin B1 to identify G2 phase cells and IdU positivity to identify S phase cells in the SPADE analysis as previously described^33^. Remaining clusters were attributed to G0/G1 phase cells (Supplementary Figure 4B).

**Figure 4:**
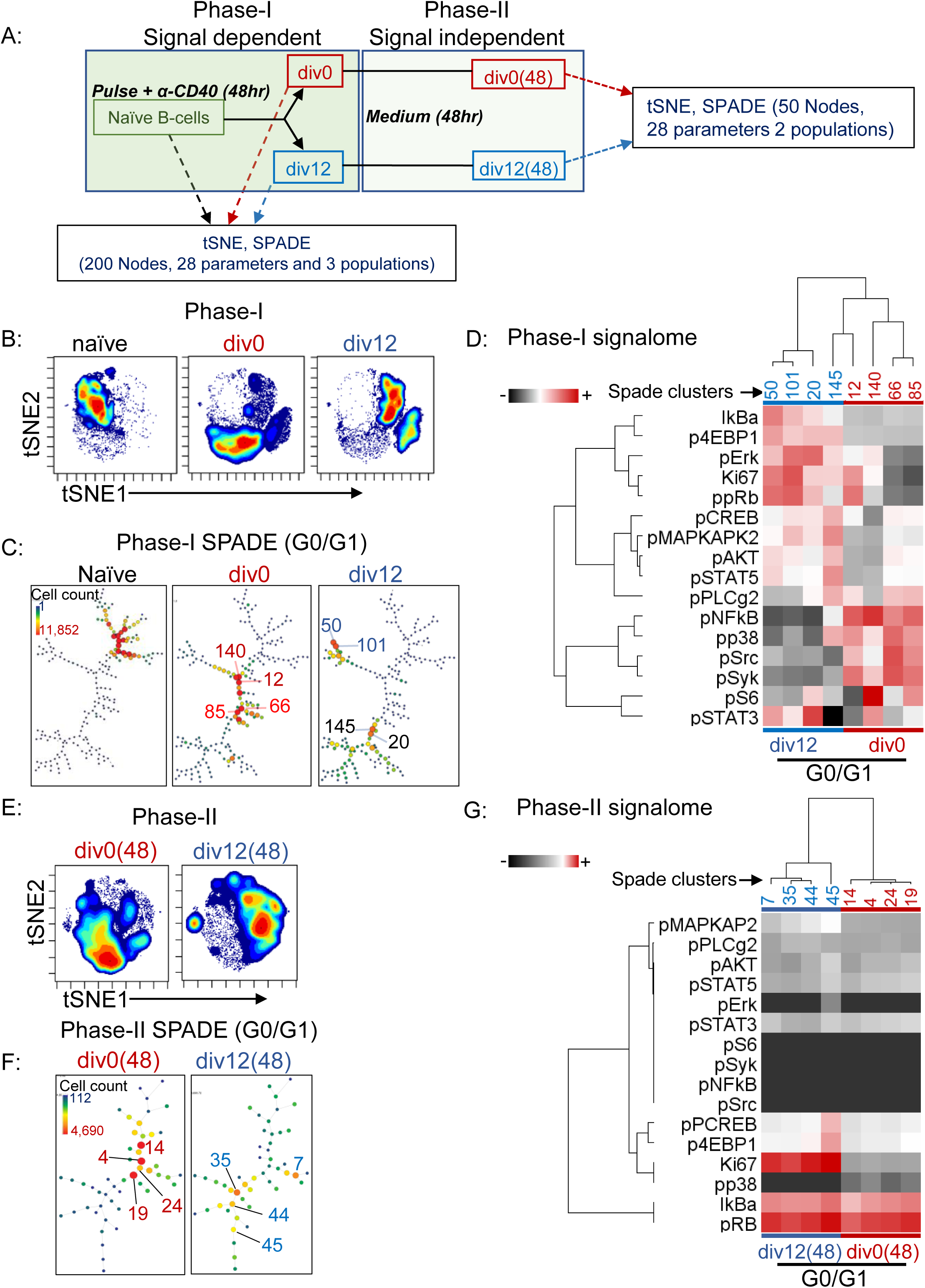
CyTOF analysis of B cell division. A) Experiment design: For analysis of signal-dependent B cell division (Phase-I), CFSE labeled B cells were activated as indicated for 48h. IdU was added to cultures for the last 30 min prior to harvest, and undivided (div0) and divided (div12) cells purified by flow cytometry. After cisplatin labeling, naïve, div0 and div12 B cells were fixed with formaldehyde and permeabilized with chilled methanol. Each population was labelled with a panel of antibodies directed against surface and cytosolic proteins (Supplementary Figure 4A). CyTOF analysis was carried out using cytobank. For analysis of cells that had undergone mitogen-independent division (Phase-II), div0 and div12 cells purified at the end of Phase-I were further cultured in the absence of mitogen for an additional 48h resulting in the cell populations labeled as div0(48) and div12(48), respectively. CyTOF analysis was carried out following IdU labeling, fixation and antibody treatments. Total 28 parameters were used to characterize 3 cell populations from Phase-I (naïve, div0 and div12) and 2 cell populations from Phase-II (div0(48) and div12(48)). B) tSNE profiles of Phase-I cell populations as indicated. C) SPADE analysis of G0/G1stage cells showing distribution of 200 nodes (n=2) in each of 3 cell populations from Phase-I. G0/G1 cells were identified by excluding S phase cells (marked by IdU) and G2/M cells marked by cyclin B and phospho-H3 (Supplementary Figure 4B). Color and size coding represent cell numbers in specific nodes. Numbered nodes were selected for further analysis (D). D) Analysis of signaling proteins in 4 SPADE nodes with the largest cell numbers from each of div0 and div12 cell populations. Heatmap representation of hierarchical cluster analysis carried out with Cluster3.0 program^55^. E) tSNE profile of Phase-II cell populations F) SPADE analysis Phase-II G1 cells showing distribution of 50 nodes (n=2) in each cell population. Cell cycle stages, colors and size of nodes are as described in part C. Numbered nodes were selected for analysis of signaling proteins in G. G) Analysis of signaling proteins in 4 SPADE nodes with the largest numbers from each of div0(48) and div12(48) cell populations. Heatmap representation of hierarchical cluster analysis as carried out in part D.

We found that G0/G1 phases of naïve, div0 and div12 cells differed greatly (Figure 4C, Supplementary Figure 4B, lower panel). Clusters of naïve cells grouped together, presumably reflecting a fairly homogenous population of G0 cells. The G1 of div0 cells was distinct from naïve B cells and more spread out across the minimum spanning tree. Two prominent groups of clusters could be discerned that likely represented different stages of G1 progression induced by continued stimulation during Phase-I. Strikingly, div12 cells segregated into two distinct, but well separated, G1 states that were distinct from the late G1 of div0 cells. Four G1 nodes each representing more than 10000 cells from div0 and div12 subsets were selected for further comparison (numbered in Figure 4C).

Each distinct G1 sub-population of div12 cells was marked by high levels of Ki67, phospho-Rb (pS807/811), phospho-Akt (pT307), phospho-Erk (pT202/pY204) and phospho-STAT5 (pY694) compared to the div0 G1 subset (Figure 4D, Supplementary Figure 4D). Additionally, div12 G1 cells also contained higher levels of phospho-4EBP1 (pT37/46), a marker for activation of cap-dependent translation. div0 G1 cells were enriched instead for signaling molecules such as phospho-Src, phospho-p38, phospho-Plcγ2, phospho-NF-κB and lower levels of IκBα relative to div12 G1 cells (Figure 4D). These observations demonstrated reduced ongoing signaling and increased expression of cell-cycle progression markers in div12 G1 cells. Conversely, div0 G1 cells had higher levels of signaling phospho-proteins. We conclude that the G1 state of cells that have divided once, differs from the state reached after persistent signaling of naïve B cells. Because many of the positive markers of div12 G1 were phosphoproteins known to be enriched in G2/M^34, 35^, we hypothesize that this state may be inherited from the preceding G2/M phase.

We also probed the intracellular states of cells that accumulated after incubating div0 and div12 cells for an additional 48h in the absence of stimulation (Figure 4A, Phase-II, div0(48) and div12(48)). The majority of cells in div0(48) comprised undivided cells, whereas the div12(48) population contained substantial numbers of cells that had undergone 1-4 mitogen-independent divisions (Supplementary Figure 4E). Accordingly, tSNE analysis of div0(48) cells showed one major cell population, whereas div12(48) cells were more heterogenous (Figure 4E). Because of reduced live cell numbers, we partitioned them into fifty clusters using the SPADE algorithm to analyze Phase-II cells (Figure 4F). We found that div0(48) and div12(48) cells had distinct G1 states (Figure 4F and Supplementary Figure 4C). In particular, div0(48) G1 cells conspicuously lacked Ki67 compared to div12(48) G1 cells despite having comparable levels of phosphorylated Rb (Figure 4G, nodes numbered in Figure 4F). A sub-population of div12(48) G1 cells were also enriched for p4EBP1, pCREB and pMAPKAPK2 compared to div0(48) G1 cells (Figure 4G). By contrast, surface receptor-initiated signaling pathways, reflected by pErk, pSrc and pSyk, were largely inactive in both div0(48) and div12(48) cells (Figure 4G). We infer that B cells can undergo several rounds of mitogen-independent cell division by propagating an activated G1 state for at least 3-4 divisions in the absence of exogenous signals.

### The post-mitotic transcriptome

To search for possible mechanisms by which such different G1 states were inherited or maintained in div0 versus div12 cells, we carried out RNA-seq from different cell subsets (Supplementary Figure 5A). We first compared the transcriptomes of purified div0 and div12 cells to their naïve precursors, Phase-I (Figure 5A). The vast majority of genes that were either up-or down-regulated in div0 or div12 cells during the first 48h culture were common to both subsets (Figure 5B). Upregulated genes shared by div0 and div12 cells were enriched for biological processes related to protein translation, mitosis/cell cycle and mitochondrial function (Supplementary Figure 5B). Down-regulated genes shared by div0 and div12 cells were enriched for processes such as immune response, adhesion and apoptosis (Supplementary Figure 5C). Smaller subsets of genes were uniquely up- (515) or down-regulated (222) in div12 cells compared to div0 cells. We hypothesized that these genes might confer the unique proliferative properties of div12 compared to div0 cells. Gene ontology (GO) analysis of div12 selective genes revealed enrichment of pathways related to centromeric chromatin and cellular metabolism (Figure 5C, Supplementary Figure 5D). Several of the chromatin associated pathways identified were related to processes that take place in G2/M and post-mitotic early G1 phases of the cell cycle^36^. These observations suggested either that 1) G2/M RNA dominated the transcriptome of div12 cells or 2) that G2/M related RNAs were present in G1 phase cells that comprised the largest subpopulation within div12. Because only 2% of div12 cells were in G2/M, we favor the interpretation that div12 G1 cells express several G2/M-selective RNAs.

**Figure 5:**
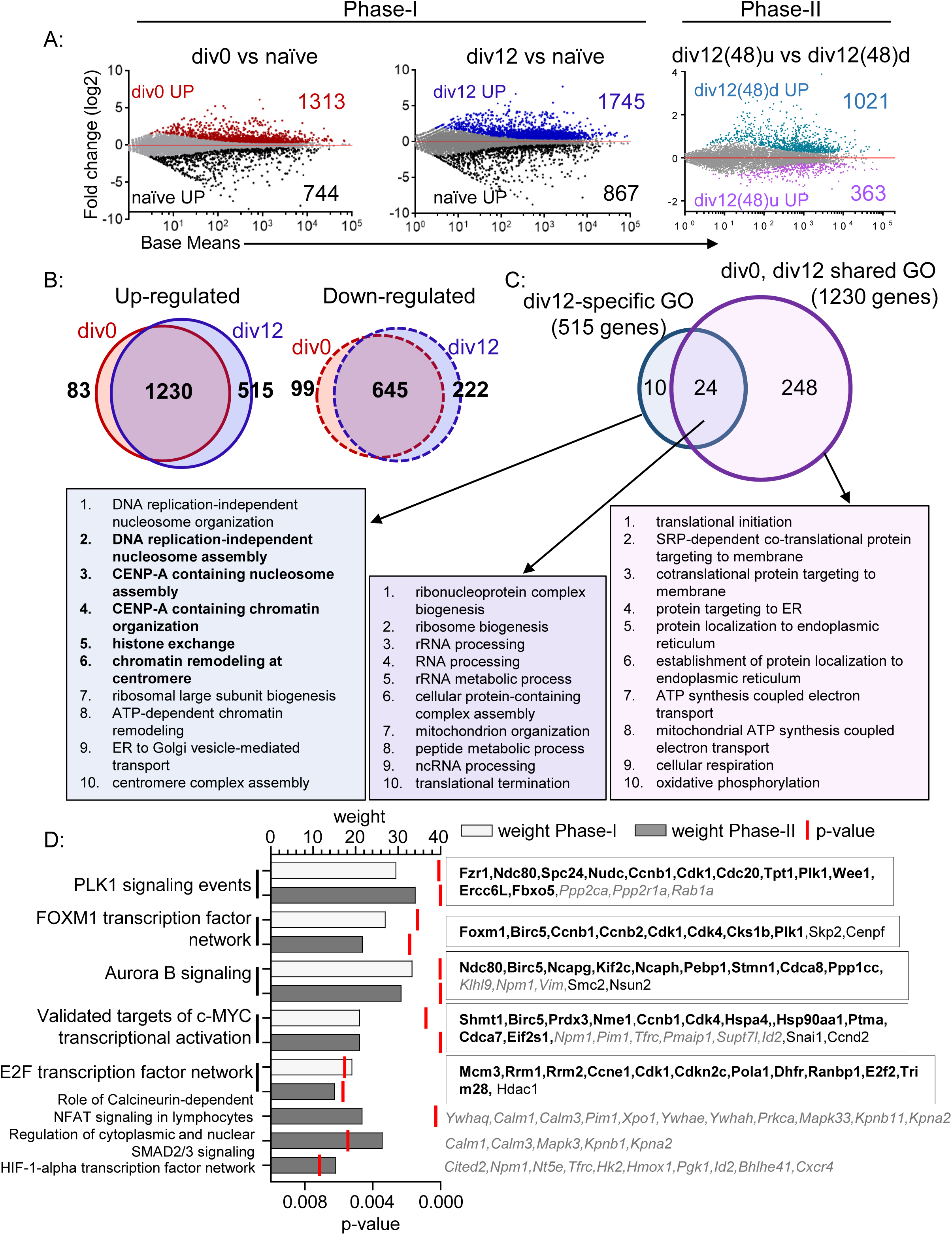
Transcriptomes of B cells undergoing mitogen-dependent and mitogen-independent division. Total RNA was prepared from 3 cell populations of mitogen-dependent Phase-I (naïve, div0 and div12) and 2 cell populations of mitogen-independent Phase-II (div12(48)d and div12(48)u corresponding to div12 cells that had divided or not during mitogen-independent culture). Following RNA-seq differentially expressed genes were identified using DESeq2 with FDR ≤0.05 for Phase-I and ≤0.1 for Phase-II. The complete experiment was repeated twice. A) MA plots showing differentially expressed genes during Phase-I and Phase-II proliferation. For Phase-I analysis gene expression was compared between div0 and naive B cells or between div12 and naïve B cells (left panels). For Phase-II analysis gene expression was compared between cells that had further divided in absence of signals (div12(48)d) and cells that had not divided further (div12(48)u) (right panel). B) Venn diagram showing overlap of up-regulated (left) and down-regulated (right) genes in div0 and div12 when compared to naïve cells. Numbers indicate genes in each category. C) Gene Ontology (GO) analysis was carried out using gene sets corresponding to up-regulated genes that were common to both div0 and div12 populations (1230 genes in part B) or genes that were specifically up-regulated in div12 cells (515 genes in part B). Venn diagram showing the distribution of GO categories in div0 and div12 cells. Top 10 GO categories (based on p≤0.05; Supplementary Figures 5B and D) are shown. D) Pathway Interaction Database (PID)^56^ analysis was carried with genes that were upregulated during Phase-I (shared between div0 and div12 cells) or upregulated during Phase-II (div12(48)d compared to div12(48)u). Enriched PIDs were scored by p-value and weight (proportion of PID genes identified in the differential gene expression analysis) (left panel). Genes shared between Phase-I and Phase-II cells are shown in bold (right panel). Genes restricted to Phase-II are shown in italics while Phase-I specific genes are non-bold.

To further identify gene expression programs of cells undergoing mitogen-independent proliferation, we compared the transcriptomes of purified cells that had or had not undergone additional cell division during 48h of Phase-II culture (Figure 5A (right), Supplementary Figure 5A). We found 1021 genes to be expressed at higher levels in cells that divided further during Phase-II. These genes were enriched for biological process such as mitotic cell cycle, protein modification and biosynthesis, indicative of their propensity to divide despite the absence of mitogen (Supplementary Figure 5F).

To search for possible gene networks that drive mitogen-dependent (Phase-I) or independent (Phase-II) cell cycle progression, Pathway Interaction Database (PID) analysis was performed. For Phase-I samples, we compared genes that were up-regulated in both div0 and div12 with naïve B cells, whereas for Phase-II analysis we compared the profiles of div12 cells that had divided during 48h culture to those that had not. PID analysis identified 5 pathways that were common to Phase-I and -II, and 3 that were unique to Phase-II (Figure 5D). All 5 shared pathways corresponded to processes involved in cell cycle progression. Of these, PLK1 signaling, the FOXM1 transcription factor network and Aurora B signaling (Supplementary Figure 5G, 5H and 5I respectively) are associated with G2/M stages of cell cycle^37, 38^. Two of the pathways uniquely present in Phase-II cells were related to calcium signaling via calmodulin and nucleo-cytoplasmic transport via karyopherin and 14-3-3 proteins. Taking a cue from the evidence of G2-like characteristics of div12 cells, we focused our attention on G2/M-specific RNAs that dominated the transcriptome of dividing B cells.

### Survivin expression in divided B cells

One G2/M specific factor that is functionally linked to centromeric chromatin (identified by GO analysis) is Birc5 (also known as survivin). Survivin is present at high levels in the G2/M phases of transformed cells, where it assists mitotic chromosome segregation via interaction with centromeric CENP proteins ^39, 40^. Pharmacologic inhibition of survivin in tumor cells results in a G2/M block followed by cell death. Survivin has also been shown to be important for G1 progression of primed CD4 T cells ^41^ and it is an integral part of FOXM1 transcription factor and Aurora B signaling networks (Supplementary Figure 5H and 5I). The predominance of CENPA-related pathways in div12 transcriptome as well as identification of the FOXM1 and Aurora B networks in cells undergoing mitogen-independent divisions prompted us to examine survivin expression in proliferating B cells.

We found that *Birc5* mRNA and protein was expressed at higher levels in div12 cells compared to div0 cells and maintained thereafter in cells that had undergone 3-4 rounds of mitogen-independent division (Figure 6A, B and Supplementary Figure 6B). *Birc5* mRNA expression pattern during Phase-I and Phase-II also correlated with *Foxm1*, *Aurkb* and *Plk1* mRNA expression (Supplementary Figure 6A and 6B). To identify the cell-cycle stage at which survivin was expressed we co-stained Phase-I cells with EdU and anti-survivin antibody for flow cytometry. As expected survivin was expressed at high levels in S and G2/M stages of both div0 and div12 cells (Figure 6C), however only div12 cells expressed survivin at the G1 stage (Figure 6C and quantified 6D). We conclude that B lymphocytes that have completed one cell division express survivin in the G1 phase. Whether this reflects retention of survivin from preceding G2/M phase or *de novo* expression in post-mitotic G1 remains to be determined. Regardless, survivin expression in div12 G1 phase imparts hallmarks of G2/M to post-mitotic G1 cells.

**Figure 6:**
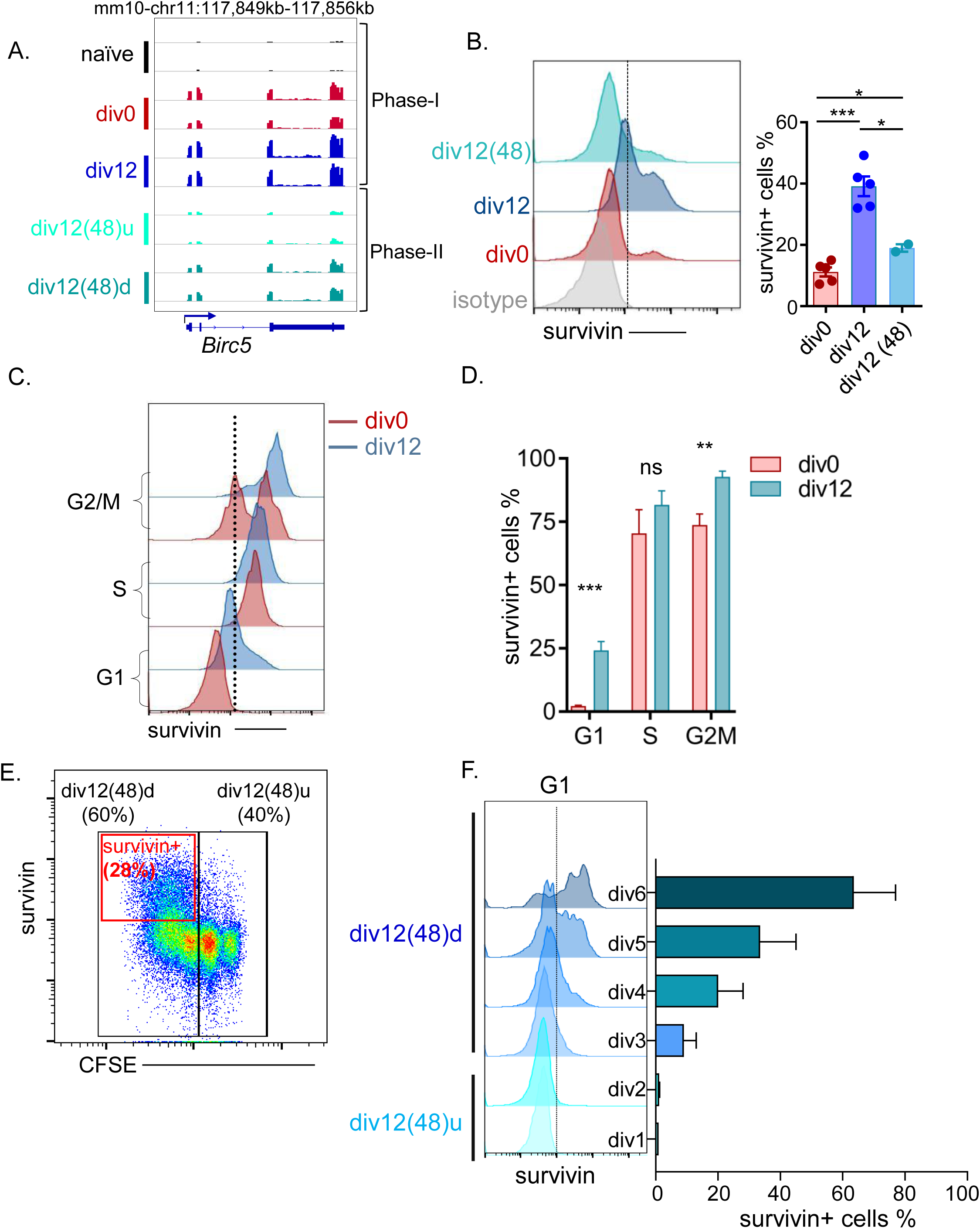
Survivin expression during mitogen-independent cell division. A) *Birc5* mRNA tracks from two biological replicates for Phase-I (mitogen-dependent) and Phase-II (mitogen-independent) cell population. div12(48)d represents cells that had undergone mitogen-independent division and div12(48)u are cells that did not further divide during the second culture. B) Survivin protein levels were assayed by flow cytometry. A representative histogram pattern is shown on the left. Div0 and div12 cells were obtained after 48h mitogenic activation as in A; div12(48) represents total cell population present after culture of div12 cells for 48h in the absence of mitogen. Right panel show the proportion of cells with survivin expression in different cell populations obtained from multiple experiments. Bars represent mean±SEM (n=5 for div0 and div12 (paired t-test) and n=2 for div12(48) (unpaired t-test)) C) Cell cycle stage-specific expression of survivin after first 48h of B cell activation as in A. Cell cycle stages were identified as in Figure 2C, using pulse EdU labeling and DAPI. A representative histogram is shown in the left panel and the average of several independent experiments shown in right panel (D). Error bars represent SEM (n=3). E) Survivin expression after 48h of mitogen-independent division of div12 cells was assayed by flow cytometry. Black boxes represent cell division states; div12(48)u cells did not divide during culture, div12(48)d cells divided during culture. Red box marks survivin positive cells. F) G1 stage-specific survivin expression in cells that have undergone different numbers of divisions from div12 cells during 48h of mitogen-independent culture. Representative histograms (left) and quantification (right; mean±SEM, n=2).

We next determined whether survivin expression in G1 persisted over the course of mitogen-independent division. For this div12 cells were cultured for an additional 48h followed by flow cytometry. We found that the cells that divided during culture expressed high levels of survivin (Figure 6E, red box). To determine the cell cycle stage at which survivin was expressed in cells that had divided in the absence of mitogen, we labelled the cells with EdU for 30min at the end of the 48h culture followed by flow cytometric analysis. We observed highest levels of survivin in the G1 stage of cells that had divided most during culture (Figure 6F). These observations demonstrate that G1-specific expression of survivin is maintained in B cells undergoing mitogen-independent proliferation.

### The role of survivin in G1-S progression in divided B cells

To evaluate the role of survivin in B cell cycle progression we inhibited its activity with the Survivin-Ran interaction blocker, LLP3 ^42^. Because effects of survivin inhibition have not been studied in primary B cells we started by assessing the effect of LLP3 on cell cycle progression of naïve B cells during Phase-I (mitogen-dependent cell division). LLP3 was added at the beginning or after 24h after initiation of B cells activation with pulse anti-IgM plus anti-CD40. Cell division and cell cycle stage were determined after an additional 30h of stimulation (Figure 7A and Supplementary Figure 7A). We found that both conditions of LLP3 treatment yielded similar results. Control cells divided 3 times during this period whereas LLP3-treated cells divided only once (Figure 7B and Supplementary Figure 7A). Importantly, S phase cells detected by a 30min pulse of EdU labeling prior to harvest were only found in the undivided population in LLP3-treated cultures. We concluded that progression from G1 to S during the first cell-cycle was not affected by LLP3, but the drug inhibited re-entry into S after the first cell division. Further characterization of cell cycle intermediates in control and LLP3-treated cells substantiated these conclusions (Figure 7C).

**Figure 7:**
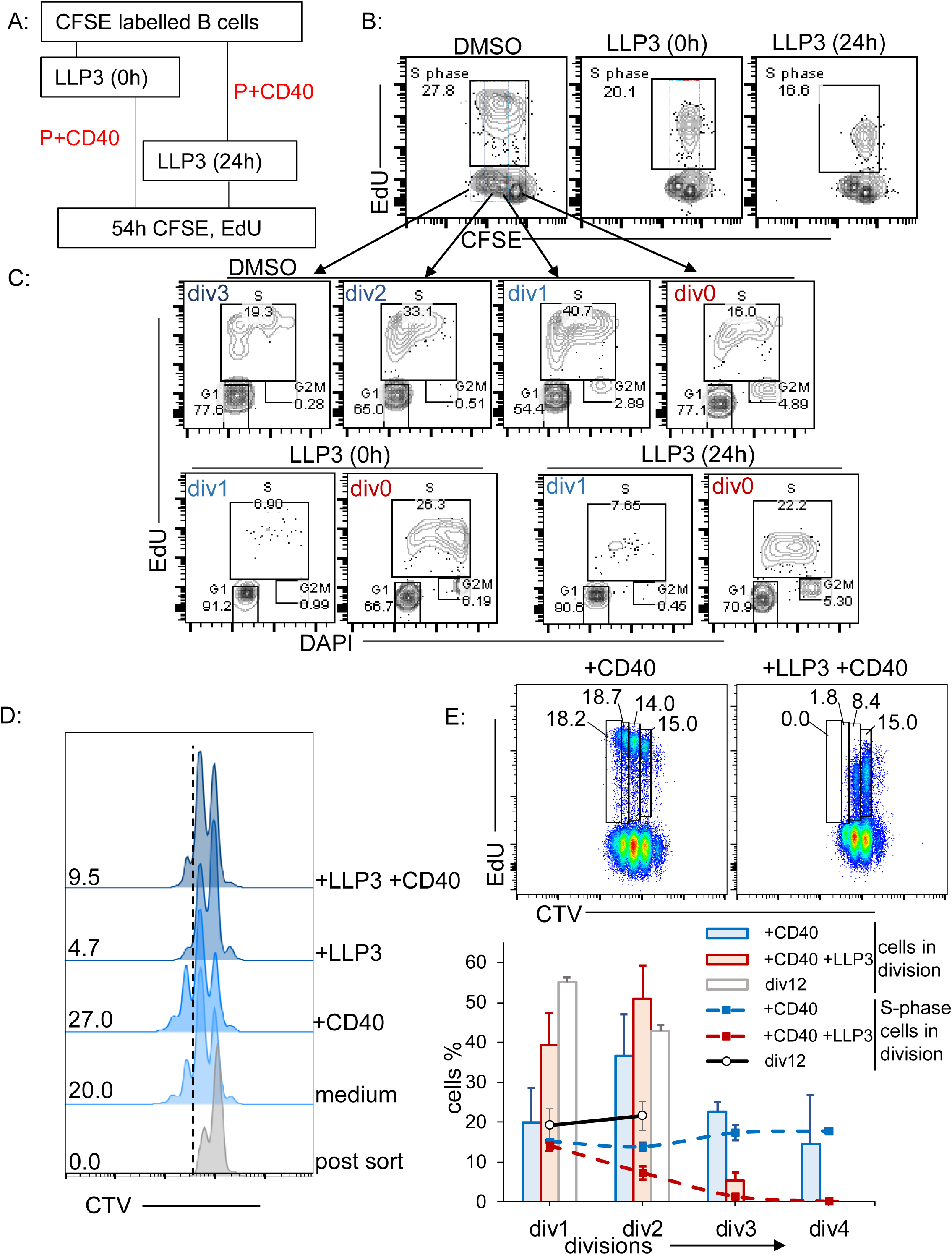
Effects of survivin inhibitor LLP3 on B cell division. LLP3 (15 µM) was added at various times during cell culture to examine the effects on mitogen-dependent (Phase-I) and mitogen-independent (Phase-II) B cell proliferation. Cell division was determined by CFSE dilution and cell-cycle stages were determined by labeling cells for 30min with EdU prior to flow cytometric analysis. A, B) Effect of LLP3 on mitogen-dependent B cell division. Schematic representation of the experimental protocol (A). B cells were activated by pulse anti-IgM and anti-CD40 in cultures that contained LLP3 from the start (0h) or where LLP3 was added after 24h. DMSO refers to cells that were not treated with LLP3. Effects on cell division were monitored at 54h, and the effects on S phase entry ascertained at the same time point by 30min EdU labeling prior to harvest (B). C) LLP3 sensitivity as a function of the number of cell divisions. Cell cycle stages of cells that had undergone 1-3 cell divisions in cultures treated with LLP3 at the beginning (0h) or after 24h as in A and B were identified by flow cytometry. D) Effect of LLP3 on mitogen-independent B cell division. Div12 cells were further cultured in medium alone or with LLP3 (15 µM), half the cultures contained low dose of anti-CD40 (0.12 µg/ml). Proliferation was assessed after 36h culture by CTV dilution. Representative histograms showing CTV dilution for indicated treatments (right), numbers within plots represent the percentage divided cells in each population. E, F) Effect of LLP3 on G1 to S progression in mitogen-independent B cells proliferation. After 36h post sort, mitogen-independent culture div12 B cells were pulse-labelled with EdU for 30min and stained using Clickit reaction before flow cytometry analysis. Representative dot-plot showing EdU-positive S phase cells and CTV dilution of cells in the presence or absence of low dose anti-CD40. Boxes and numbers represent percentage of S phase cells in each division (E). Two independent experiments from part E were quantified. Bars show the percentage of cells present in each division after 36h of mitogen-independent culture. Lines represent percentage of cells present in S phase of each division. Error bars represent SEM. Composition of div12 cells post-sort are shown in grey bars and corresponding S phase cells in black lines.

We next assessed the requirement for survivin during mitogen-independent proliferation. For this purified div12 cells obtained after Phase-I were re-cultured in the presence or absence of LLP3 for an additional 36h. Cell division was measured by CTV dilution. Cells that had divided 2 or 3 times amounted to approximately 20-30% of the culture after 36h in the absence of LLP3. Inclusion of LLP3 during secondary culture reduced mitogen-independent proliferation 3-4-fold regardless of the presence of survival signals provided by low dose of anti-CD40 (Figure 7D). To directly assay S phase cells, we included EdU during the last hour of secondary culture. EdU-positive cells were markedly reduced in the small number of cells that newly emerged in LLP3 treated cultures (Figure 7E). We conclude that survivin is an essential mediator of G1 to S progression during mitogen-independent proliferation.

## Discussion

Primary splenic B lymphocytes are present in the G0 cell cycle state prior to antigen encounter. The first critical event during a humoral response is clonal expansion of a tiny fraction of cells that recognize the newly encountered antigen via their cell surface antibody receptors. This initial expansion increases the numbers of antigen-specific cells by several orders of magnitude. In current models the extent of lymphocyte proliferation has been linked to the concept of division destiny, a cell autonomous condition induced during primary antigenic encounter^15^. Division destiny is proposed to be the cumulative outcome of multiple signals that impinge on cells responding to antigen. During T-independent responses BCR signaling alone is a major driver of clonal expansion, whereas additional signals provided by T cells determine the outcome of T-dependent responses. Mechanisms that drive multiple rounds of cell division during clonal expansion remain unclear. One possibility is that persistence of antigen permits re-stimulation of proliferating B cells after each division. Alternatively, cellular states established before the first mitosis may permit several rounds of cell division in the absence of continued re-stimulation. The former model is consistent with prevailing views of cell cycle regulation that posit recurrent mitogenic signaling being required for G1 progression after each cell division. By contrast, several studies, especially in CD8^+^T lymphocytes^20, 21^, indicate that the cell cycle may be uniquely configured in lymphocytes that allows the second form of mitogen-independent proliferation. However, mechanisms that permit lymphocytes to circumvent the classical model remain obscure.

Here we show that the G1 state of B lymphocytes after the first cell division is distinct from the G1 state of activated B cells that have not divided. This state is characterized by the presence of hyper-phosphorylated Rb, phosphorylated-CDK2 and phosphorylated-Akt (T308). These are hallmarks of G2/M cell cycle stages that are believed to be inactivated by phosphatases in late G2^34^. Additionally, hallmarks of G2/M were also evident in RNA-seq analysis (see below). We propose that, contrary to prevailing dogma, B cells carry over hallmarks of G2 into the post-mitotic G1 phase. This state permits G1 progression without requiring renewed mitogenic stimulation. Unlike observations in some tumor cells where persistent CDK2 activity was shown to circumvent the need for CDK4/6 activation for renewed G1 progression, mitogen-independent G1 progression of B cells requires CDK4/6 activity. Importantly, B cells undergoing mitogen-independent proliferation maintained their size over several divisions, likely due to enhanced metabolic activity despite the absence of mitogen (AS, data not shown). This condition was distinct from division destiny induced by persistent TLR9 stimulation during which cells reached a non-proliferative, quiescence state characterized by reduced cell size^15, 43^. During the time course of our studies, mitogen-independent proliferation of B cells was limited only by accentuated cell death which could be attenuated via non-mitogenic CD40 stimulation. We propose that the ability of B cells to undergo multiple cell divisions with limited stimulation is an important determinant of clonal expansion.

These observations raised two related questions. First, how do features of G2/M phases persist in the post-mitotic G1 phase and second, what drives continued mitogen-independent cell division? With regards to the first question, the prevailing view is PP1 and PP2A activity in late G2/M de-phosphorylates key proteins, such as pRb, p130^34^ and AKT^35^ and also inactivate Ras signaling^44^, thereby resetting the clock in the newly generated G1. In contrast, we found that several of these PP1 and PP2A target proteins remained hyper-phosphorylated in the G1 of B cells that had divided once. We infer that phosphatase activity is attenuated in G2/M of primary B lymphocytes, thereby leading to inheritance of a partially activated G1 after cell division. Reduced PP2A activity may also underline elevation of the Ca++/calcineurin pathway in post-mitotic cells noted in our PID analysis (ref fig). We also observed elevated levels of Myc in post-mitotic G1 phase, though we cannot distinguish whether Myc levels were also inherited from the preceding G2/M stages or synthesized *de novo* in the new G1 stage. Our transcriptomic analyses revealed up-regulation of validated targets of Myc during mitogen-independent cell division, supporting a role for Myc. Because ectopic Myc expression is not sufficient to induce cell division in B cells^45^, we hypothesized that persistent Myc expression contributes to mitogen-independent cell division by maintaining an elevated metabolic state via expression of its target genes^45^.

In order to understand mechanisms that fueled persistent cell division, we compared the metabolic state of B cells prior to, or just after, first cell division. We found that basal oxygen consumption as well as peak mitochondrial respiration were increased in post-mitotic cells (data not shown). Attempts to evaluate the role of PI3K using pharmacologic inhibitors were unsuccessful due to excessive cell death in cultures treated with inhibitors. Interestingly, mitogen-independent cell division was affected minimally by inclusion of rapamycin in post-mitotic cultures suggesting that mTOR-dependent pathways were not involved in the process. However, elevated cell death noted in such cultures preclude definitive conclusions about the role of mTOR in this process.

Though we found that CDK4/6 activity was necessary for G1 progression during mitogen-independent division, a key aspect that remains unclear is the identity of cyclins that activate CDK4/6. Our transcriptomic studies indicated that levels of cyclin D2 mRNA were low in cells that have divided once, however, we could not directly evaluate protein levels by flow cytometry. Recent studies using ES cells triple-deficient for D-type cyclins suggest that such cyclins may not be essential for G1 progression^23^. It is possible that CDK4/6 activity in these cells is maintained by such a cyclin-independent mechanism.

Comparing transcriptomic signatures and annotated protein interaction database of cells that were permissive for mitogen-independent proliferation, we identified a link to genes relevant to G2/M phases of the cell cycle. Amongst these genes, *Birc5* (survivin) caught our attention as one with prior connection to multiple stages of the cell cycle regulation. Though primarily known as a factor important for chromosome segregation and anti-apoptotic during mitosis in tumor cells^46^, survivin has also been proposed to activate CDK4/6^47^ and for G1 progression of primed CD4+ T cells^41^. We found that survivin expression in G1 stage cells correlated with the propensity to undergo mitogen-independent cell division. Indeed, survivin expression that was induced transcriptionally during the first G1 was maintained, and even increased, in cells that had divided multiple times without exogenous signals. A functional role for survivin was further demonstrated by modulating survivin function using LLP3. Though the first G1 progression of resting B cells in response to mitogenic signals was minimally affected by LLP3, we found that passage through subsequent G1 states to S phase was significantly adversely affected. Importantly, mitogen-independent cell division was reduced 4-fold in the presence of LLP3. We propose that survivin is an essential component of the program that permits mitogen-independent G1 progression after the first cell division in B cells.

In summary, we provide evidence that primary B cells can undergo several rounds of cell division without requiring a G1 progression signal after each mitosis. We hypothesize that this ability is important for clonal expansion during a primary immune response. It also provides a plausible explanation for a puzzling feature of germinal centers (GC) where T-dependent B cell signals have been shown to occur in the GC light zone whereas the bulk of B cell division occurs in the dark zone, apparently in the absence of cell stimulation. Thus, requirements for B cell proliferation are distinct from current views of the cell cycle and more in line with properties of tumor cells, a proportion of which have been shown to be capable of G1 progression in the absence of growth factor signals. However, primary B cell undergoing such mitogen-independent proliferation lack survival signals that are a hallmark of transformed cells. It is likely that microenvironmental signals in secondary lymphoid organs cooperate with this cell cycle property of B cells to tune the extent of expansion during immune responses.

## Author contributions

A.S. and R.S. designed research; A.S., J.P.J., M.K. and X.Q. performed research experiments; A.S., M.H.S. and G.P.N. performed CyTOF experiments and analysis; A.S. and R.S. analyzed data; A.S. and R.S. prepared manuscript.

## Acknowledgements

We thank all the members of Sen Lab in Gene Regulation Section (GRS) for valuable discussion. This work was supported in part by the Intramural Research Program of the NIH, National Institute on Aging (NIA).

## Methods

### Mice

All experimental mice used were 8–12 weeks old C57BL/6J mice purchased from The Jackson Laboratory and maintained in house facility at NIA, Baltimore.

Mice were treated humanely in accordance with federal government guidelines. The protocol was approved by the Animal Care and Use Committee of the NIA Intramural Research Program,

### B cell isolation and activation culture

Primary B cells were isolated using EasySep B cell kit (StemCell Technologies) according to manufacturer’s protocol and were ≥95% pure based on CD19 staining. Purified B cells (1.5×10^6^/ml) were cultured in c-RPMI (RPMI 1640 (Invitrogen) supplemented with 10% heat-inactivated FBS (Gemini), 55mM β-mercaptoethanol (Sigma), 2mM L-glutamine and 100-IU penicillin and 100 μg/ml streptomycin (Gibco)) at humidified 37°C incubator. Phase-I cells were cultured in 100mm tissue culture dishes (Corning) while Phase-II cells were cultured in 24 well tissue culture plates (Corning).

For proliferation assays, B cells were labeled with cell trace dyes either Carboxyfluorescein succinimidyl ester (CFSE) or Cell Trace Violet (CTV) (Invitrogen) according to manufacturer protocol. These labeled or non-labeled cells were pulsed or continuous activated with goat anti-mouse IgM F(ab’)2 (Jackson Immunoresearch) as described earlier^31^. After 6h hamster anti-mouse CD40 monoclonal antibody (BD Pharmingen) was added and proliferation was measured by dilution of cell trace dye at different time points by flow cytometry. For other mitogenic stimulations, CD40 alone, 5 μg/ml anti-mouse CD40 monoclonal antibody (BD Pharmingen) while LPS (Sigma) 5 μg/ml or CpG 50 ng/ml (Invivogen) was added after CFSE or CTV labelling for indicated times.

### Flow cytometry

For non-fixed cells live cells were identified using either TOPO3 (Invitrogen) or DAPI (Invitrogen) negative staining. Fixed live cells were identified by either forward and side scatter gates or by using fixable viability dye (FVD780) (BD Biosciences). Cells were counted using TC20 (Biorad) or Precision Counting Beads (Biolegend) according to manufacturer’s protocol. Divided and non-divided B cells were sorted based on cell trace dyes levels using FACS Aria II or FACS Aria Fusion (BD Biosciences). Sort sensitivity was set to fine-tune and post sort purity was consistently ≥95%. For Phase-II, cells were washed with warm complete RPMI after sorting, counted and plated (1.5×10^6^/ml) in c-RPMI at humidified 37°C incubator for 1h to recover and then indicated treatments were performed.

### Antibodies and reagents

Complete list of antibodies related to activation, flow cytometry, western blotting and CyTOF are listed in Supplementary Table 1 with section a, b, c and d respectively. Inhibitors and other reagents are listed in Supplementary Table 2.

### Cell cycle and intracellular staining analysis

Proliferating cells were pulse labelled for 30 minutes with EdU (Invitrogen) and Clickit reaction was performed according to manufacturer’s protocol to label incorporated EdU with Alexa Flour dyes. For flow cytometry analysis of intracellular proteins cells were fixed using 1.6% methanol free paraformaldehyde (Electron Microscopy Science, EMS) for 15min at room temperature and washed twice with PBS and resuspended in ice cold chilled 500 µl permwash buffer III (BD Biosciences) and incubated for at least 30 minutes at 4°C or stored in −20°C till next step. After two washes cells were resuspended in 100µl saponin buffer (Invitrogen) and primary antibody was added with optimal concentration and incubated at 4°C for 1h. For non-conjugated primary antibodies, fluorochrome labeled secondary antibody were used according to manufacturer’s protocol. For cell cycle analysis using DNA content, fixed cells were incubated for 30 min at 37°C in staining buffer prepared of saponin buffer (Invitrogen) containing DAPI (1 µg/ml) (Sigma) and RNAse (10 µg/ml) (Invitorgen).

For CyTOF staining, proliferating B cell cultures were pulsed for 30 min labelled with IdU (10 µM) (Sigma) for S phase cell identification at indicated time points. After harvesting at indicated time, for dead cell staining, cultured B cells were resuspended in 5ml of Cisplatin (50 µM) (Sigma) in PBS for precisely 60s and quenched immediately with 2X volume of complete RPMI. Cells were further washed 2x with stain buffer (BD Bioscience) and fixed with 1.6% final concentration of methanol free paraformaldehyde for 15 minutes at room temperature. After washing cell pellets were stored in 2% BSA in PBS at −20°C till staining with surface and intracellular antibodies was performed according to previously described method^48^.

All measurements for flow cytometry were carried out using FACS CantoII or FACS Aria Fusion (BD Biosciences) and CyTOF acquisition was performed on CyTOF^TM^ mass cytometer (DVS Sciences, Toronto, Canada). Data was analyzed using Cytobank (tSNE and SPADE) (https://cytobank.org/) or FlowJo software (Tree Star).

### Western blotting

Whole cell extracts (WCE) were made using RIPA buffer (Thermo Scientific) supplemented with protease and phosphatase inhibitor cocktail (Thermo). Proteins were separated by electrophoresis through 10-20% SDS-PAGE and electrophoretically transferred to nitrocellulose membrane using iBLOT2 (Thermo). After blocking with 5% BSA in Tris HCl-0.05% Tween (TBST) for 1h at room temperature, membranes were incubated with primary antibodies overnight at 4°C, washed in TBST, and incubated for 1h with horseradish peroxidase (HRP)–coupled secondary antibodies (Cell Signaling). Blots were developed using the enhanced chemiluminescence (ECL) systems (Pierce) and captured by GeneSys Syngene Imaging System.

### RNA-seq and gene expression analysis

Two independent biological replicates each consisting pools of 4-5 mice were used for Phase-I RNA-studies (Supplementary Figure 5A). For Phase-II, divided (div12) B cells were further cultured in presence of 0.12 µg/ml of anti-CD40 for next 48h and further FACS sorted in divided (div12(48)d) and non-divided (div12(48)u) cells. Total RNA from 1×106 (Phase-I) and 5×105 (Phase-II) cells was isolated using Qiazol (Qiagen) according to manufacturer’s protocol. RNA-sequencing of total RNA was performed at the Johns Hopkins Deep Sequencing and Microarray Core facility using standard protocol for NextSeq 500 sequencer. Briefly, ribosomal RNA depleted was depleted from 100ng of total RNA and barcoded library was made and 50 bp single end reads were generated from each sample. Reads were analyzed using online server Galaxy RNA-seq pipeline (usegalaxy.org). Adapter trimmed sequences were aligned to mouse genome (mm10) using HISAT2^49^. Estimation of transcripts was done using Salmon^50^ algorithm (TPM) and featureCounts^51^ and htseq-count^52^ were used for differential gene expression analysis using DESeq2^53^. Gene ontologies and PID analysis were performed using ToppFun^54^ suite online server at ToppGene (https://toppgene.cchmc.org/). For PID weight score calculation, following equation was used:

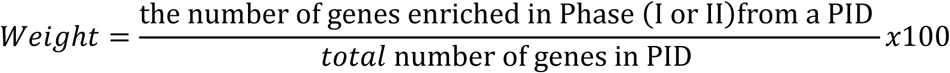

### Inhibitor experiments

Two independent biological replicates each consisting pools of 2-3 mice were used for CDK inhibitor experiments. After sorting cells from Phase-I into divided (div12) and non-divided (div0) B cells were cultured in presence of various CDK inhibitors or DMSO for Phase-II (next 48h).

During Phase-II divided div12 B cells cultures were having only medium while div0 cells were supplemented with anti-IgM (10 µg/ml) for re-stimulation. Inhibitors and their doses are listed in Supplementary Table 2. Proliferation was measured by CFSE dilution and cell cycle was analyzed using DNA content analysis by DAPI staining.

For survivin inhibition experiments two independent biological replicates each consisting pools of 3 mice were used. Survivin inhibitor, LLP3 (15 µg/ml) (Sigma) dissolved in DMSO was added at indicated times.

### Statistical analysis and graphs

Statistical tests and graphs were created using Prism (version 7.0) (GraphPad) or Excel (Microsoft). Data were considered statistically significant at *p ≤ 0.05, **p ≤ 0.01 and ***p ≤ 0.001.

