## Supplementary figures for "Mitogen-independent cell cycle progression in B lymphocytes"

**Supplementary Figures 1-7**

Mitogen-independent cell cycle progression in B lymphocytes.

**Singh et al.**

Supplementary Figure 1

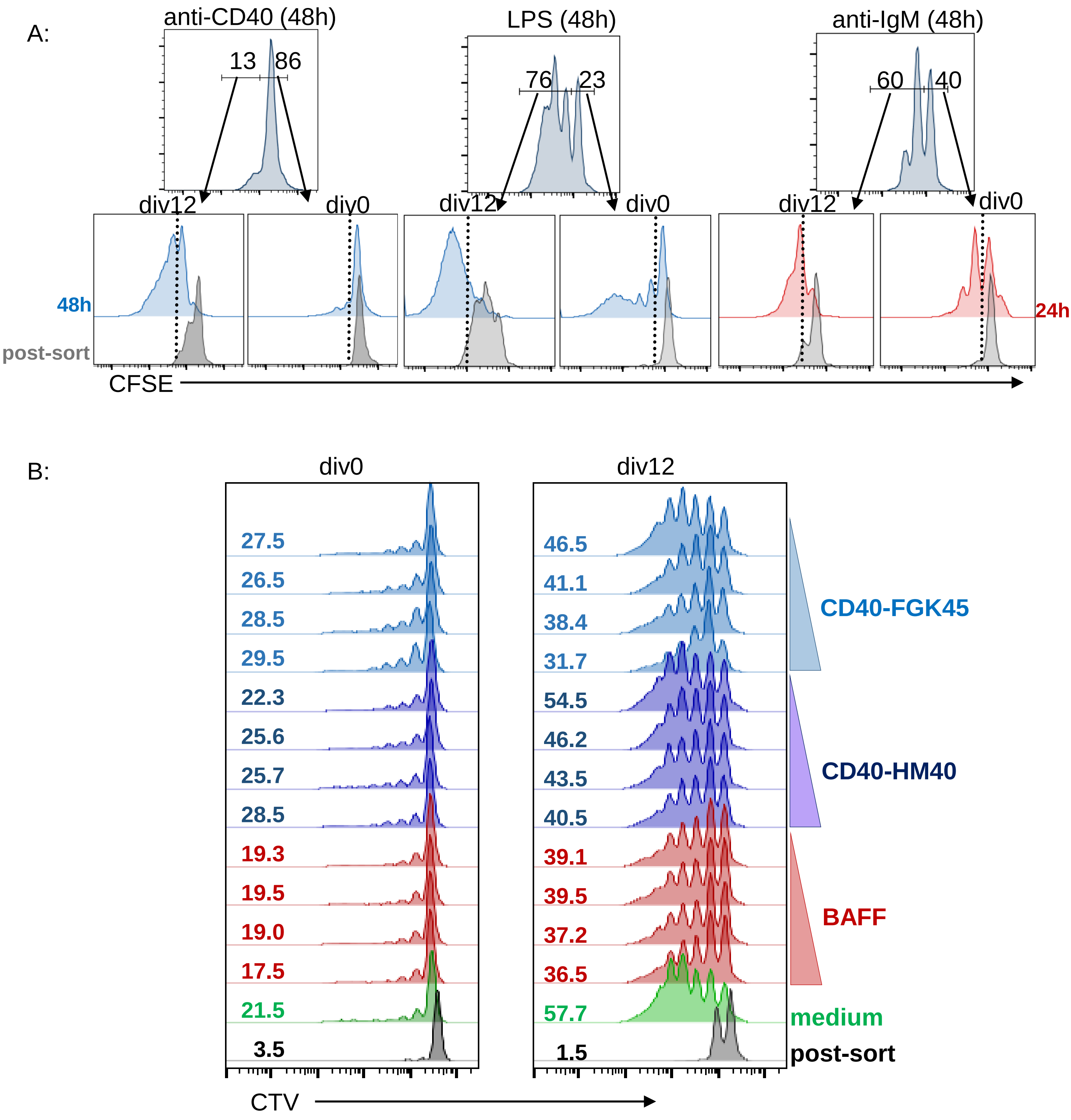

C:

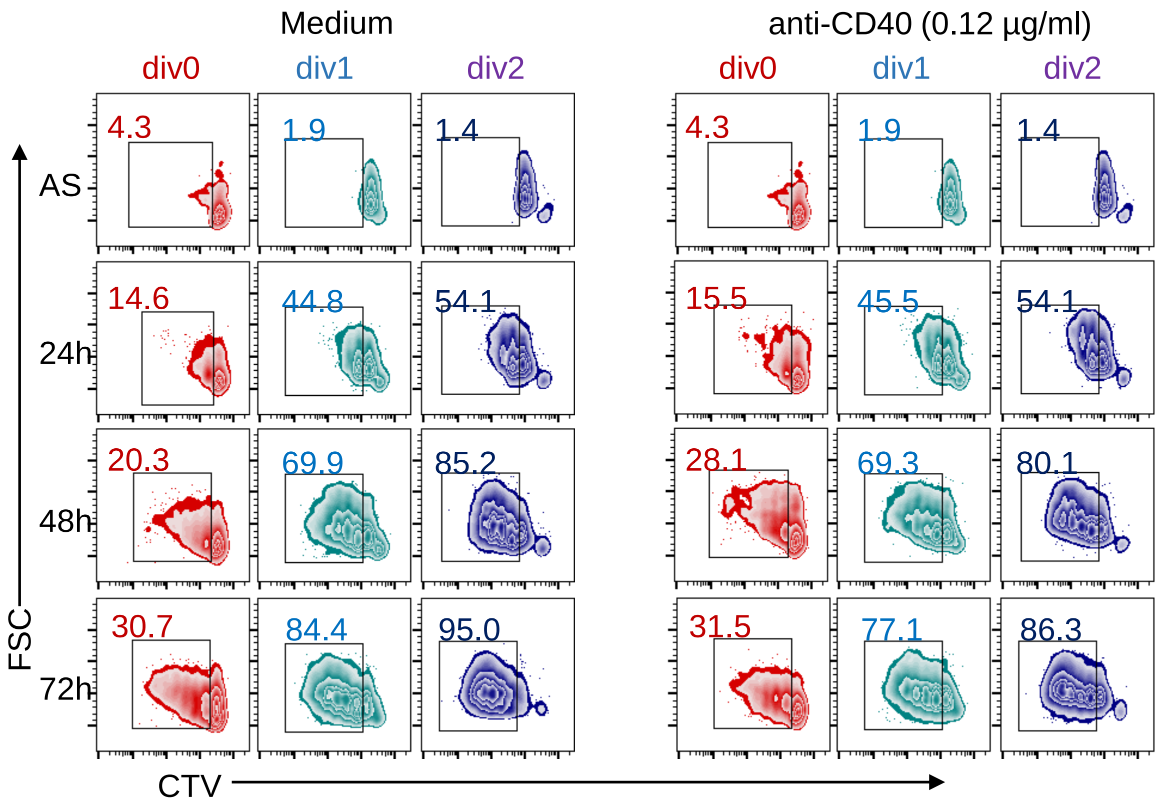

Medium
anti-CD40 (0.12 µg/ml)
div0
div1
div2
div0
div1
div2
AS
24h
48h
72h
FSC
CTV
4.3
1.9
1.4
4.3
1.9
1.4
14.6
44.8
54.1
15.5
45.5
54.1
20.3
69.9
85.2
28.1
69.3
80.1
30.7
84.4
95.0
31.5
77.1
86.3

Supplementary Figure 2

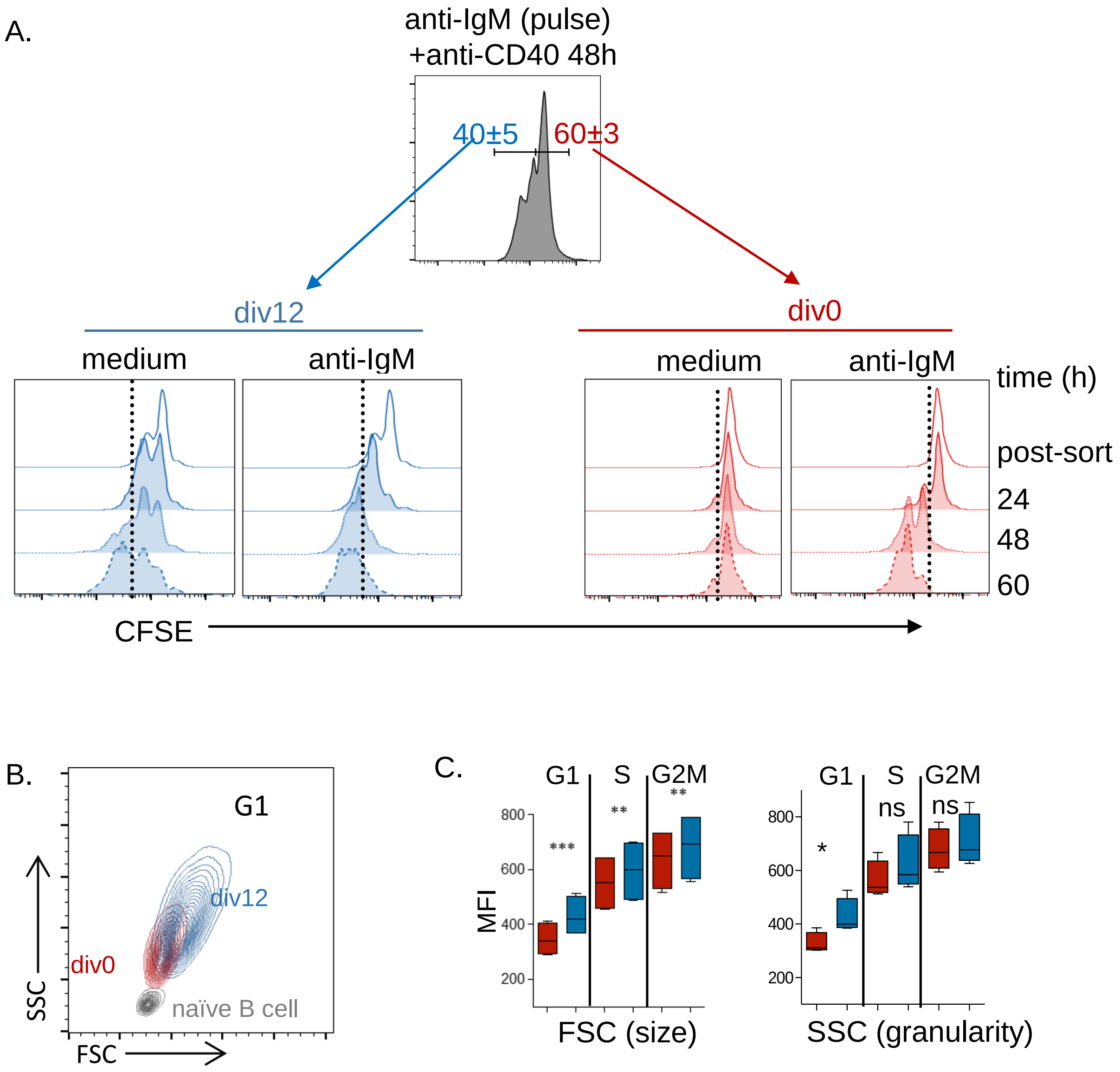

Supplementary Figure 4

A:

| Gating (9) | Cell cycle (5) | Signaling (18) | | Others (7) |
|---|---|---|---|---|
| *CD19* | *pRb* | *p4EBP1* | *pCREB* | *IgM* |
| *B220* | *CyclinB1* | *IkBa* | *pPLCg2* | *IgD* |
| CD3 | *IdU* | *pSrc* | *pSTAT5* | *MHCII* |
| Cisplatin | *Ki67* | *pp38* | *pSTAT1* | *CD23* |
| DNA1 | *pHH3* | *pSyk* | *pNFkB* | *CD27* |
| DNA2 | | *pErk* | *cPARP* | CD45 |
| Event Length | | *pAkt* | *pS6* | CFSE for phase II |
| Time | | *pSTAT3* | | |
| Bead Density | | *pMAPKAPK2* | | |

B:

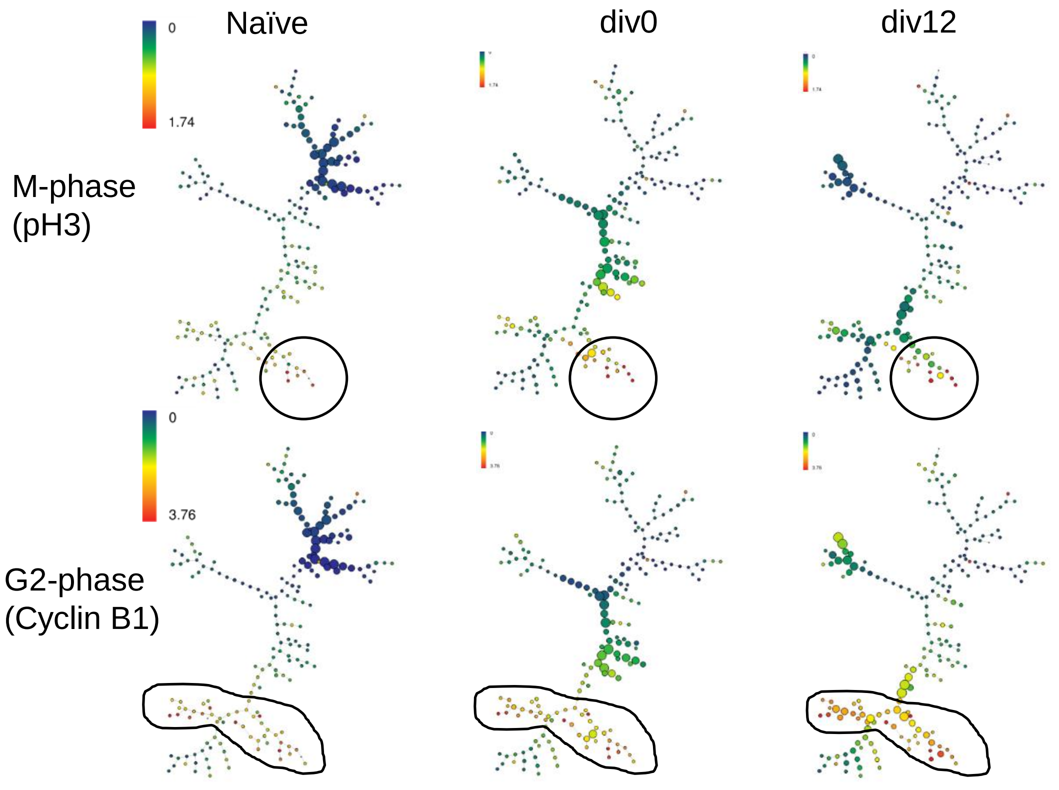

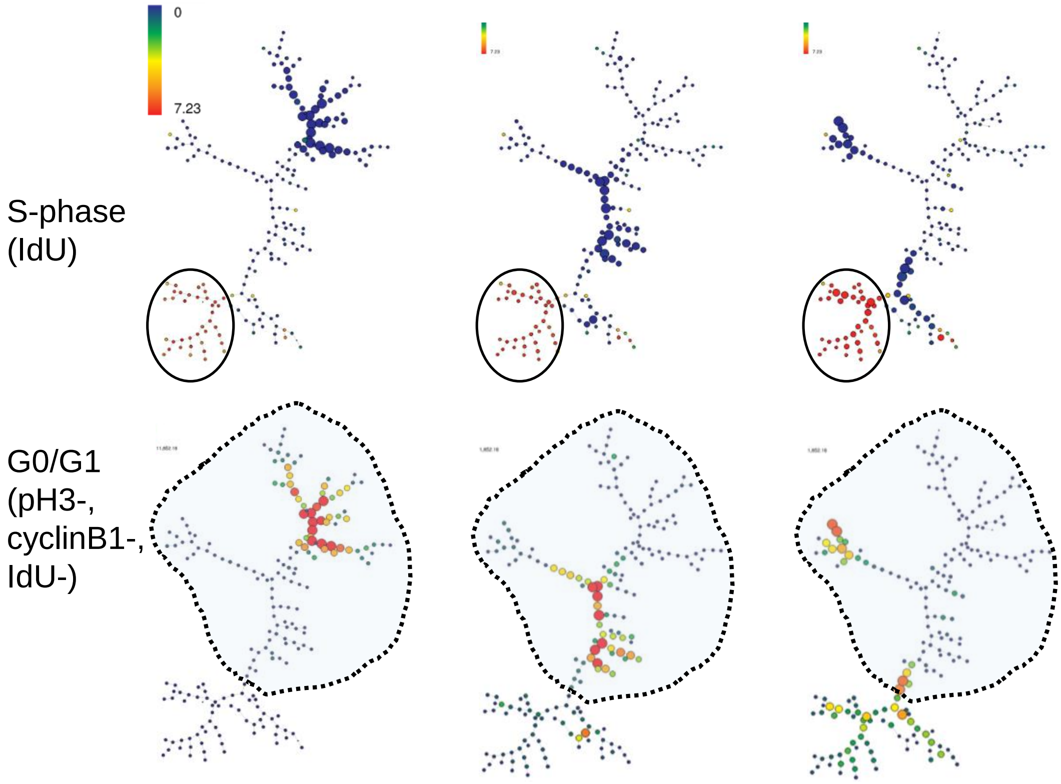
0
7.23
S-phase
(IdU)
G0/G1
(pH3-,
cyclinB1-,
IdU-)

C:

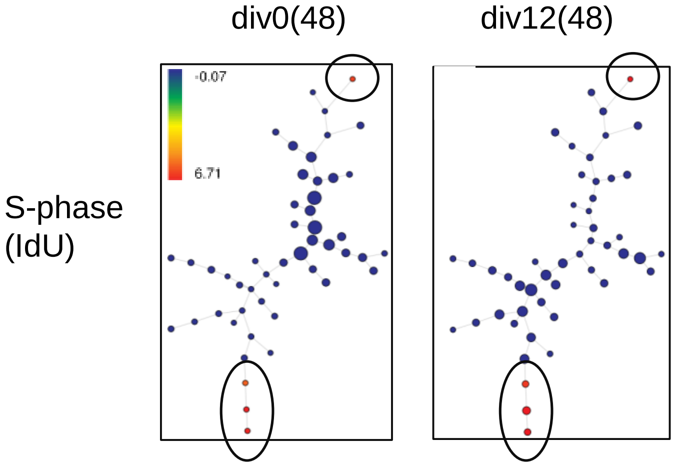
div0(48)
div12(48)
-0.07
6.71
S-phase
(IdU)

D:

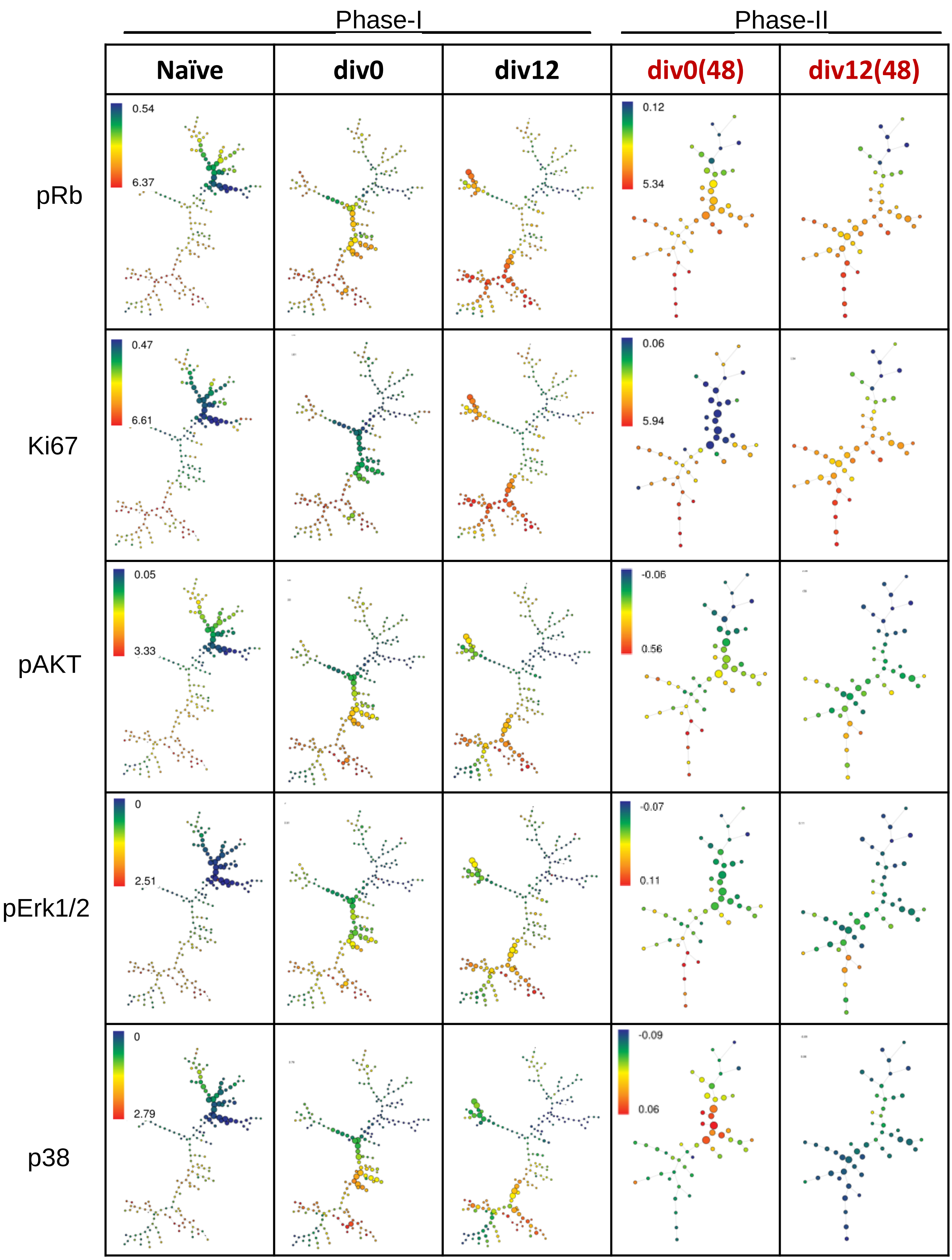
Phase-I
Phase-II
Naïve
div0
div12
div0(48)
div12(48)
pRb
0.54
6.37
0.12
5.34
Ki67
0.47
6.61
0.06
5.94
pAKT
0.05
3.33
-0.06
0.56
pErk1/2
0
2.51
-0.07
0.11
p38
0
2.79
-0.09
0.06

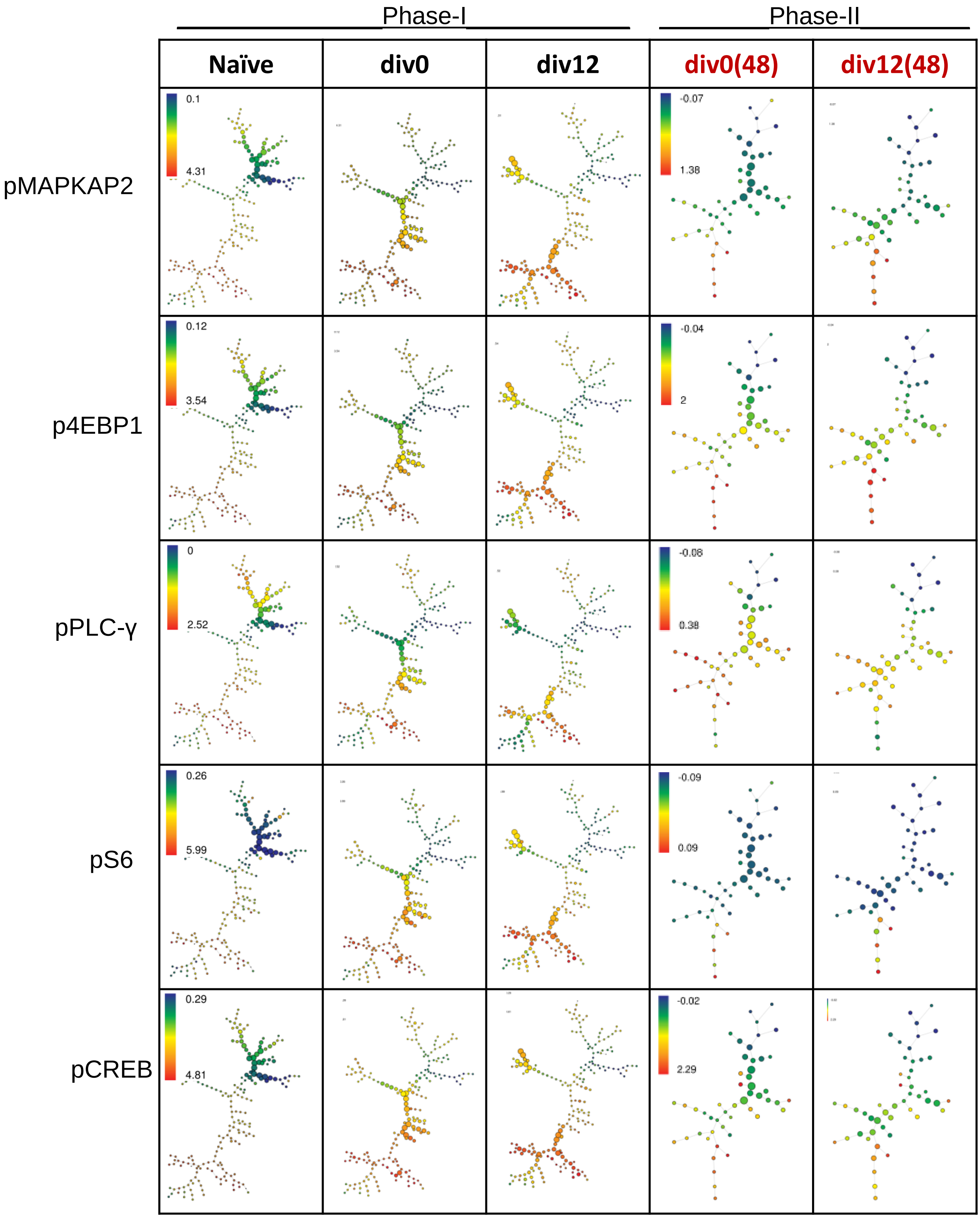
Phase-I
Phase-II
Naïve
div0
div12
div0(48)
div12(48)
pMAPKAP2
0.1
4.31
-0.07
1.38
p4EBP1
0.12
3.54
-0.04
2
pPLC-γ
0
2.52
-0.08
0.38
pS6
0.26
5.99
-0.09
0.09
pCREB
0.29
4.81
-0.02
2.29

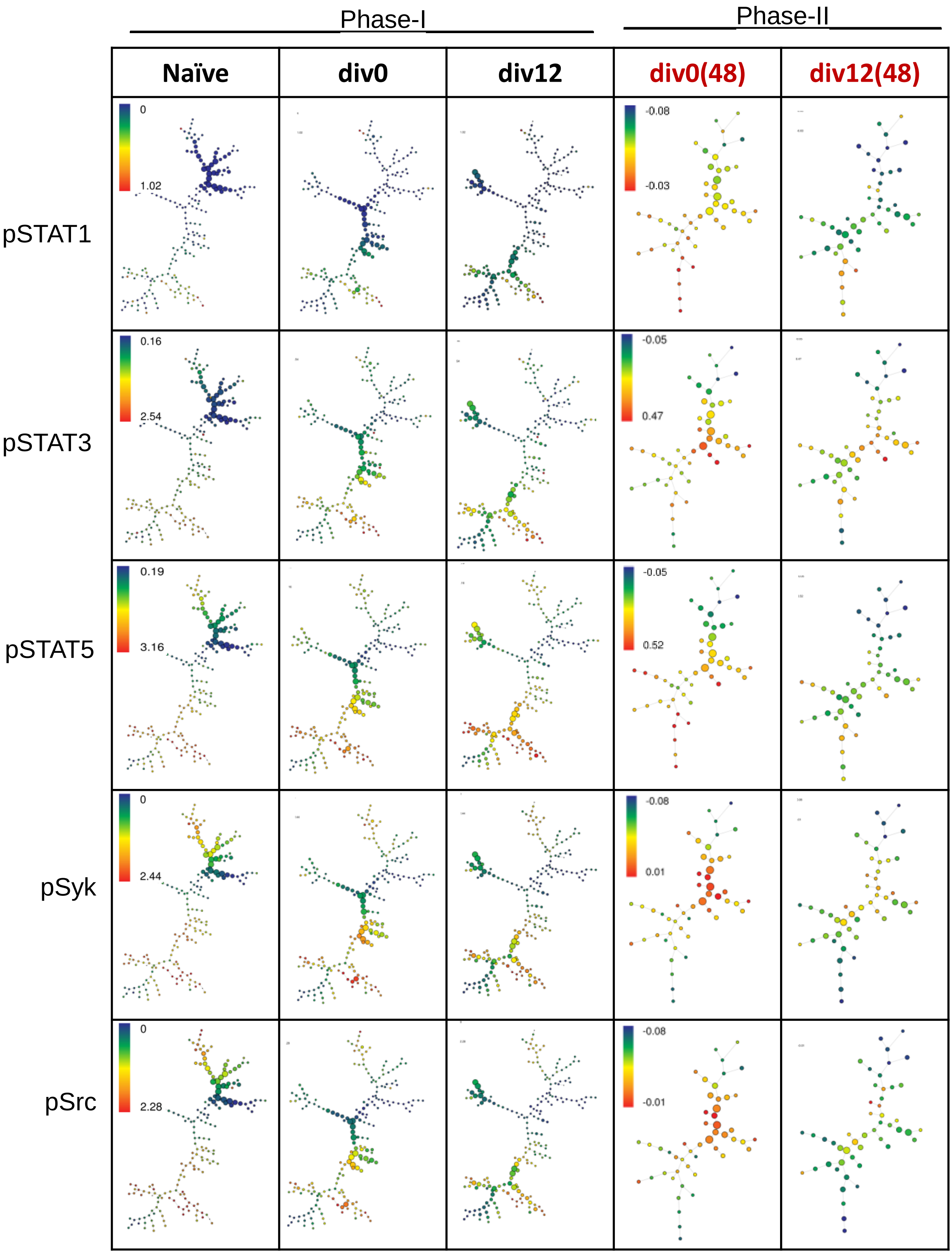

Phase-I
Phase-II
Naïve
div0
div12
div0(48)
div12(48)
pSTAT1
pSTAT3
pSTAT5
pSyk
pSrc
0
1.02
-0.08
-0.03
0.16
2.54
-0.05
0.47
0.19
3.16
-0.05
0.52
0
2.44
-0.08
0.01
0
2.28
-0.08
-0.01

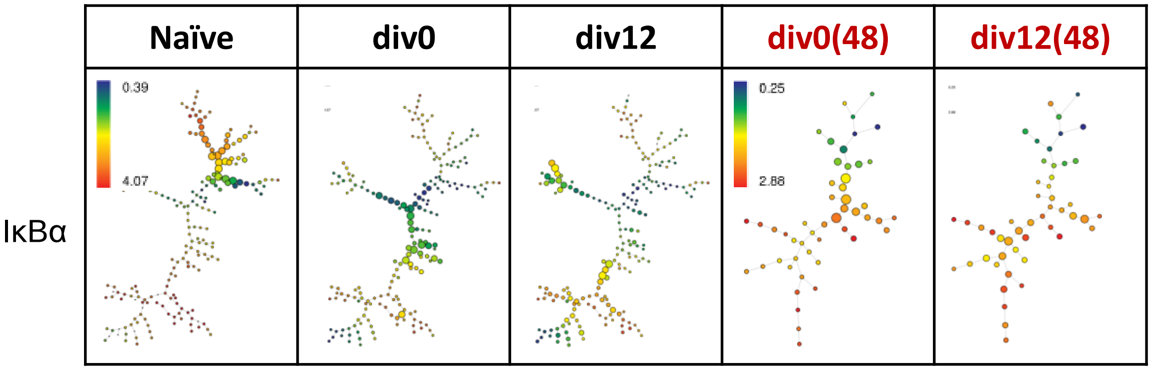
Naïve
div0
div12
div0(48)
div12(48)
IκBα
0.39
4.07
0.25
2.88

E:

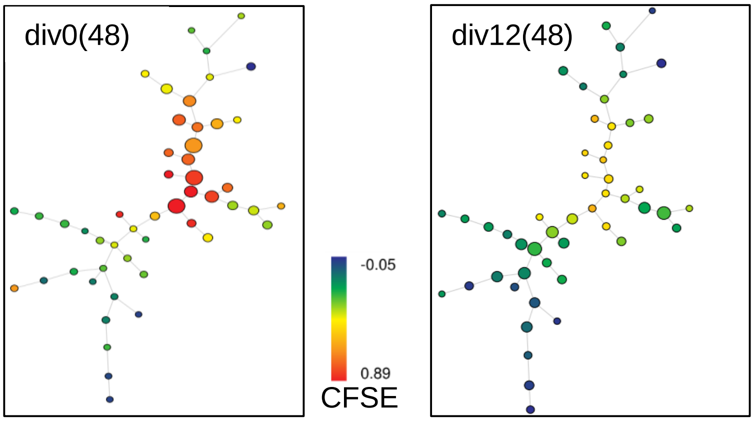
div0(48)
div12(48)
-0.05
0.89
CFSE

Supplementary Figure 5

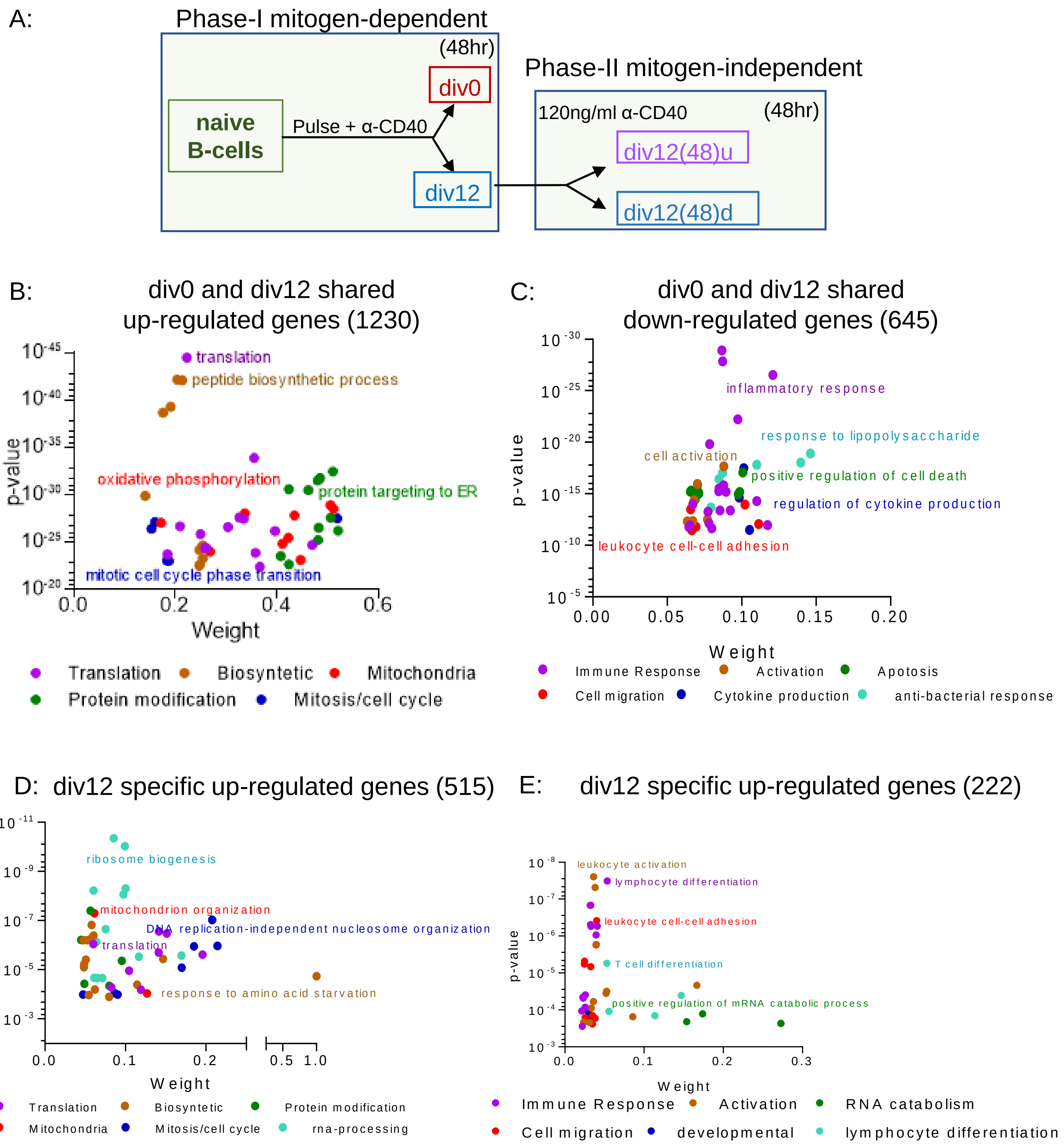

F:

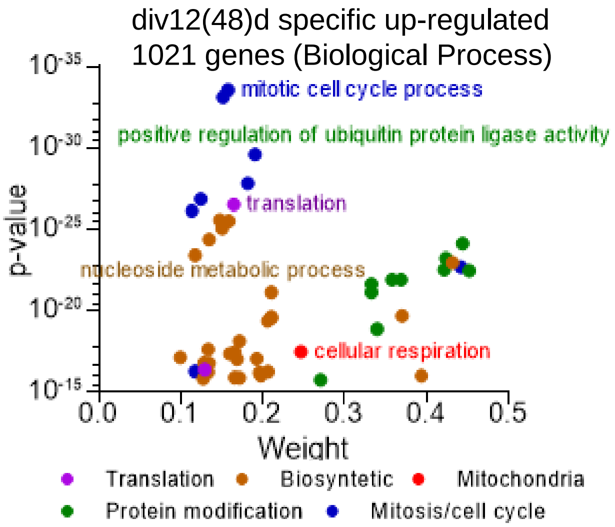


G: PLK1 signaling events

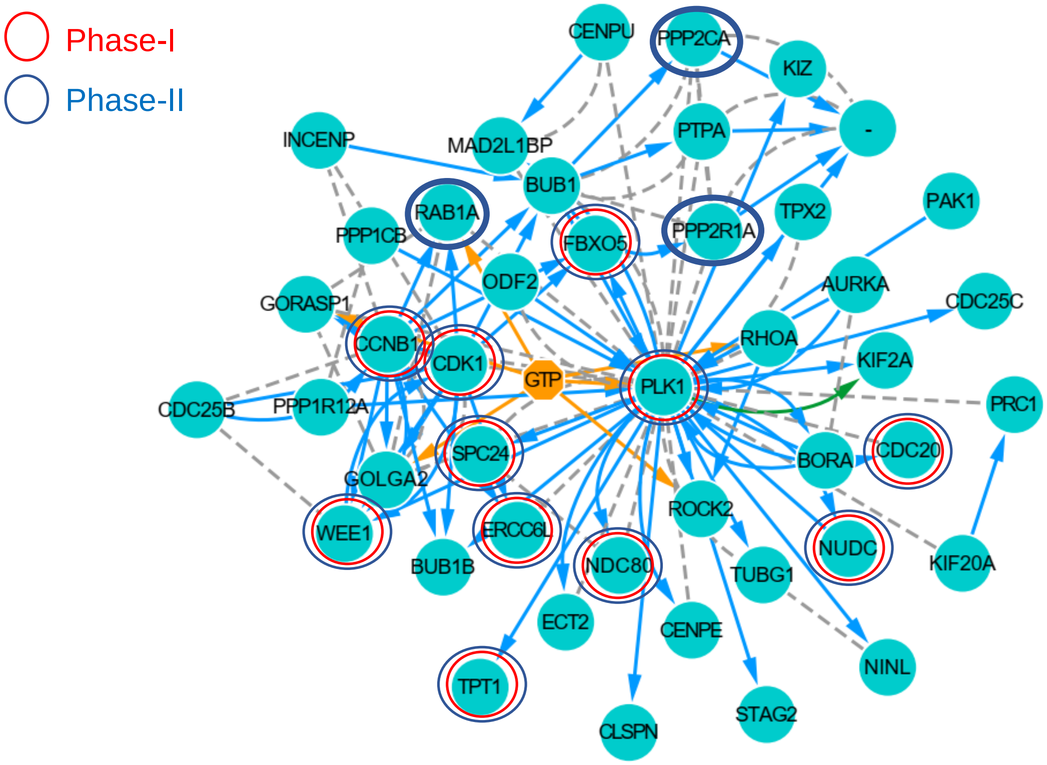

H: FOXM1 Transcription Network

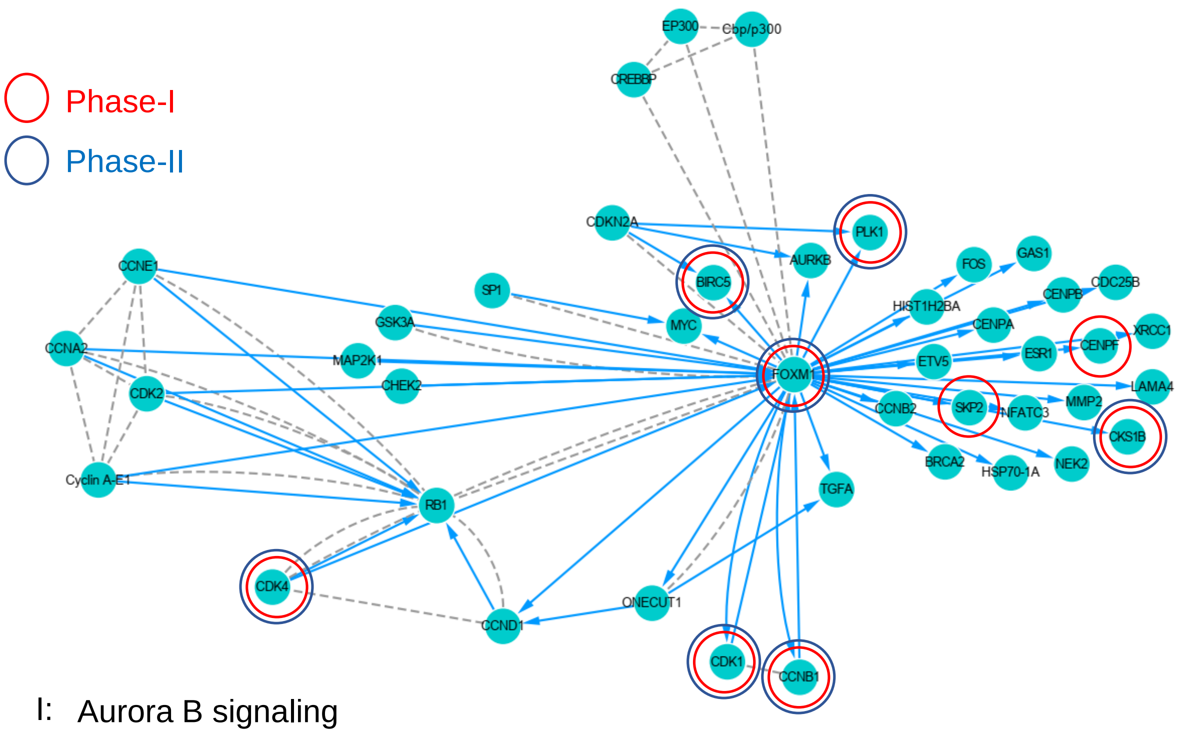


I: Aurora B signaling

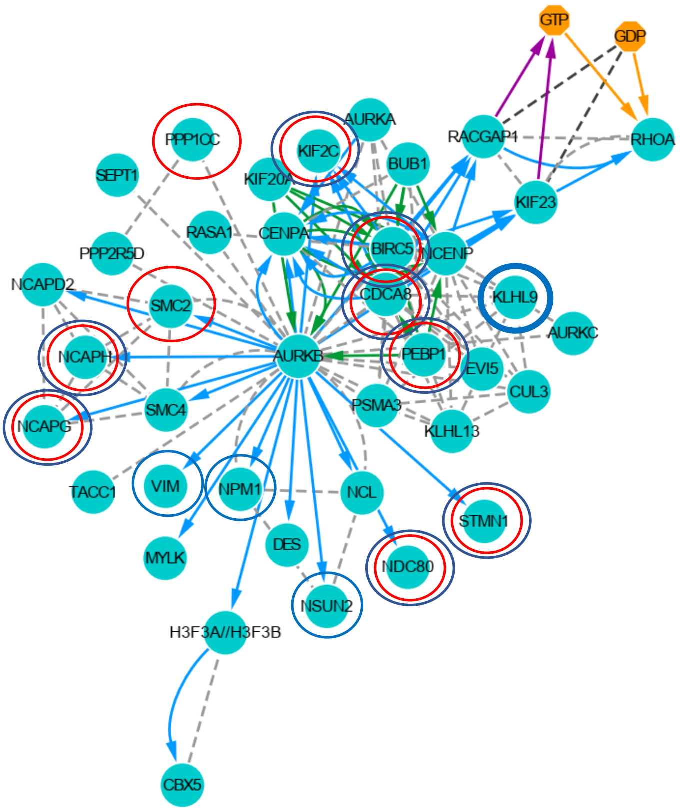

Supplementary Figure 6

A:

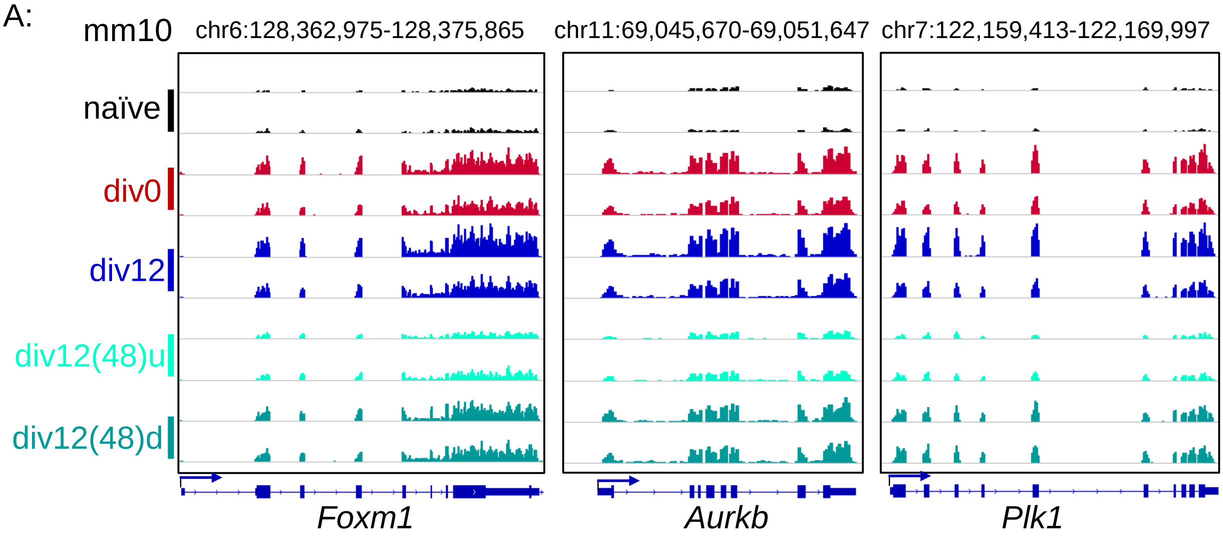


B:

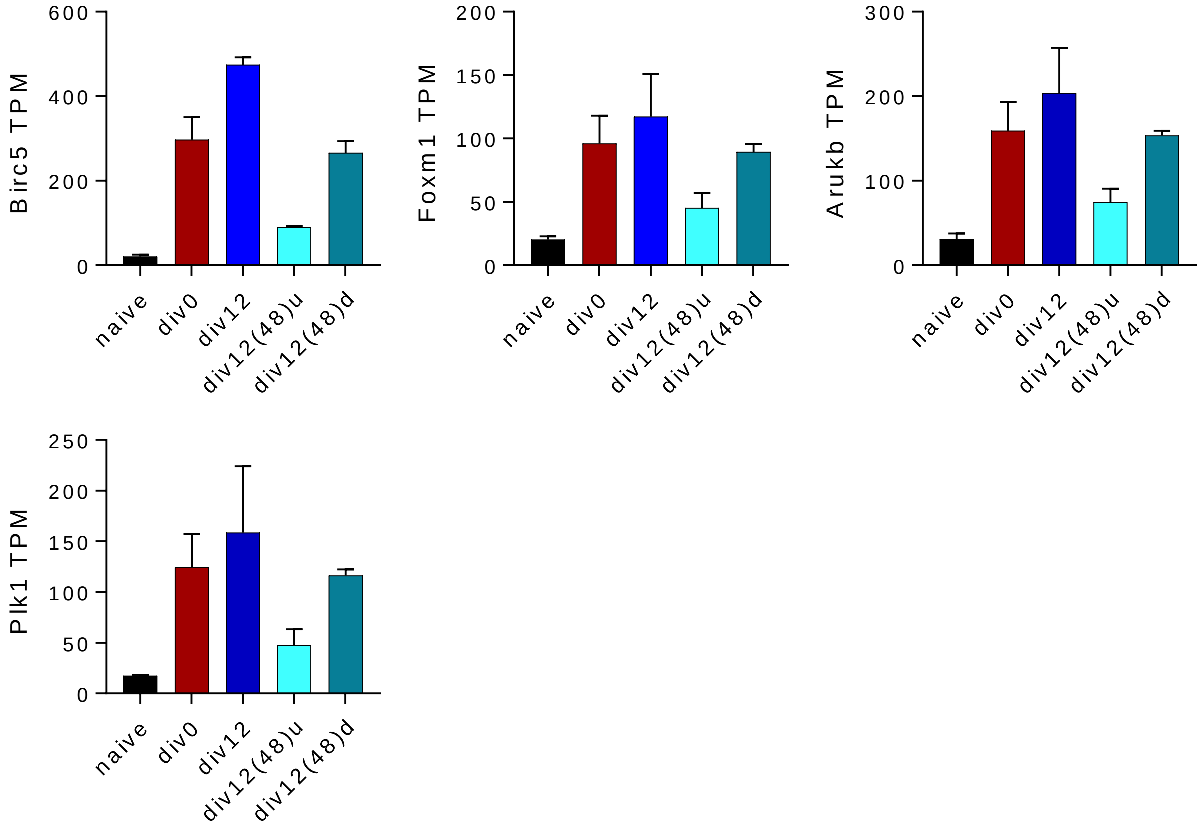

Supplementary Figure 7

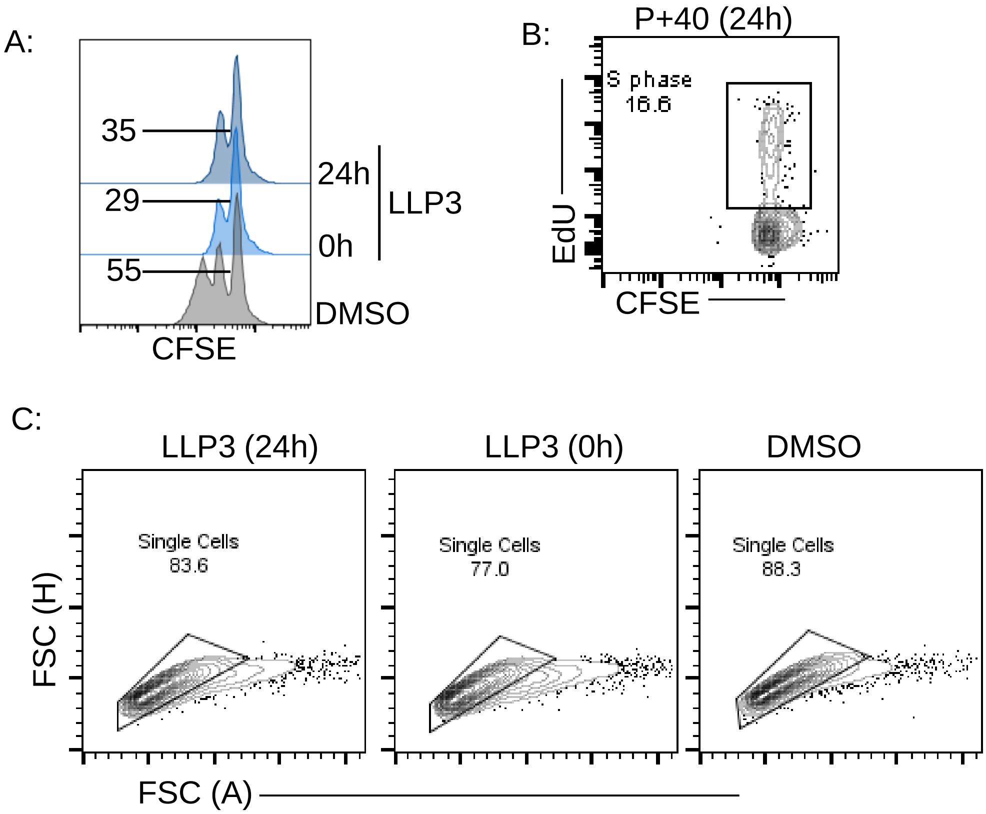
