## Supplementary figure legends for "Mitogen-independent cell cycle progression in B lymphocytes"

Supplementary Figure 1:

Effect of different mitogenic stimulus leading to first mitotic division and subsequently leading to mitogen-independent B cell proliferation. CFSE labelled splenic B cells were activated for 48h with indicated mitogens (anti-CD40 (5 µg/ml), LPS (5 mg/ml) and anti-IgM Fab’2 (10 µg/ml)). Divided (div12) and non-divided (div0) B cells were sorted using FACS and further cultured without any mitogen for next 48h. Proliferation was readout as CFSE dilution measured using flow cytometry.

A) Representative histograms of mitogen-dependent proliferation profile of indicated mitogen at 48h (top). Gates within represent the divided and non-divided B cells. CFSE profiles after post-sort and 48h later for anti-CD40 or LPS activated and post-sort and 24h for anti-IgM activated B cells (bottom). Dotted horizontal line represents distinction between divided and non-divided B cells.
B) Influence of survival signals from BAFF and anti-CD40 antibody on mitogen independent B cell proliferation. div0 and div12 B cells were sorted as Figure 1C, and further cultured for 48h without any stimulus (medium) or varying dose of BAFF (100-10 ng) or two different clones of anti-CD40 antibody (5-0.12 µg/ml). Histograms representing CTV dilution at 48h post-sort of div0 (left) and div12 (right) with indicated stimulus on right (bold letters). Numbers represent the percentage of divided B cells during 48h after sorting.
C) Change in size (forward scatter, “FSC”) during mitogen-independent B cell proliferation. Zebra-plot showing FSC and CTV dilution for no-stimulation (left, “medium”) or anti-

CD40 (right, “anti-CD40 0.12 μg/ml”) after 72h post sort from B cells isolated as Figure 1C. Numbers represent the percentage of divided B cells.

Supplementary Figure 2:

A) Non-divided B cells are not refractory to proliferation. Divided (div12) and non-divided (div0) B cells were sorted as Figure 1C and cultured further with (anti-IgM) or without stimulus (medium) for 60h. Representative histogram representing CFSE dilution with indicated time post-sort. Horizontal dotted line indicates distinction between divided and non-divided B cells.

B) Activated yet not divided (div0) cells are bigger than their precursor naive B cells and divided B cells (div12) are even bigger than non-divided (div0). Contour plot representing forward scatter (size, FSC) and side scatter (granularity, SSC) from the G1 phase of naïve (resting), activated but non-divided div0 or divided div12.

C) Quantification of part B for div0 and div12 from 5 independent experiments. Forward scatter (left) and side scatter (right) are shown by box and whisker plot demonstrating the mean fluoresce intensity (MFI) and error bars SEM with paired t-test. ***=p<0.0001, **=p<0.001, *=p<0.05 and ns=not significant.

Supplementary Figure 4:

A) Parameters used during CyTOF analysis. Parameters in italics were used in tSNE and SPADE analysis.

B) Gating strategy on SPADE plots for identification of various stages of cell cycles. First row, mitotic cell (M-phase) nodes identified by high levels of pH3, second row, G2 phase cell nodes identified by high levels of Cyclin B1 and third row, shows S phase cell nodes as identified by high levels of IdU. Absence of all these marks are demonstrated in forth row as G0/G1 cells.

C) Nodes in SPADE plots representing S phase cells during mitogen-independent proliferation if Phase-II for div0 (left) and div12 (right).

D) SPADE plots for indicated markers during Phase-I (left 3 columns) and Phase-II (right 2 columns)

E) SPADE plots representing CFSE levels during Phase-II mitogen-independent proliferation.

Supplementary Figure 5

A) Schema of RNA seq experiment, 3 populations form Phase-I (naïve, div0 and div12) and 2 populations from Phase-II (div12(48)u and div12(48)d) were analyzed for their transcriptome analysis using RNA-seq.

B-F) Scatter plots representing enriched Biological Process identified by GO analysis using TOPP gene for indicated comparisons. Plots demonstrated p-value and weight of corresponding biological process. Similar biological process are represented by same color and representative process is shown on the graphs.

G-I) Pathway Interaction Database (PID) networks for indicated network. Phase-I nodes are circled red while Phase-II by blue color.

Supplementary Figure 6

A) Foxm1, Aurkb and Plk1 mRNA tracks from two biological replicates from indicated cell populations.

B) Bar graphs representing TPM values from two biological replicates calculated using Salmon algorithm. Cell population are on X axis while Y axis represent TPM value for indicated mRNA.

Supplementary Figure 7

A) Histograms demonstrating effects of pharmacological inhibition of survivin by LLP3. CFSE dilution was accessed at indicated 54h with indicated time treatment of LLP3 cell culture as shown in Figure 7 A.

B) Representative contour plot showing EdU positive cells (representing S phase cells) and CFSE levels in pulse anti-IgM+ anti-CD40 (P+40) treated cells at after 24h.

C) Representative contour plot showing presence of single cells in P+40 activated cells at 54h treated with DMSO at 0h or LLP3 at indicated times. Single cells were identified by plotting forward scatter area (FSC (A)) vs forward scatter height (FSC (H)), single cells populate at the diagonal in these plot and are represented by polygonal gates
