## Supplementary tables for "Mitogen-independent cell cycle progression in B lymphocytes"

| Supplementary Table 1 | | | | | |
|---|---|---|---|---|---|
| **a (activation)** | | | | | |
| **Target Protein** | **Clone** | **Vendor** | **Concentration (ug/ml)** | | |
| AffiniPure F(ab')$_2$ Fragment Goat Anti-Mouse IgM, µ chain specific | Polyclonal | Jackson Immunoresearch | varies (5-10ug/ml) | | |
| anti-CD40 | HM-40 | BD | varies (125ng/ml-5ug/ml) | | |
| anti-CD40 | FGK | Novus Biologicals | varies (125ng/ml-5ug/ml) | | |
| **b (flow cytometry)** | | | | | |
| **Target Protein** | **Clone** | **Vendor** | **Concentration/Dilution** | | |
| CD19 | 6D5 | Biolegend | 1:200 | | |
| pRb (S807/811) | D20B12 | CST | 1:500 | | |
| cMyc | D84C12 | CST | 1:200 | | |
| pCDK2 (Thr160) | Thr160 | CST | 1:50 | | |
| Survivin | 71G4B7 | CST | 1:500 | | |
| Rabbit IgG (isotype) | DA1E | CST | 1:500 | | |
| anti-rabbit IgG Fab2 PE | | CST | 1:1000 | | |
| anti-rabbit IgG Fab2 AF-647 | | CST | 1:500 | | |
| **c (western blotting)** | | | | | |
| **Target Protein** | **Clone** | **Vendor** | **Concentration (ug/ml)** | | |
| p27 | SX53G8.5 | CST | 1:1000 | | |
| b-actin | AC-74 | Sigma | 1:5000 | | |
| anti-rabbit IgG-HRP | | SANTA CRUZ BIOTECHNOLOGY | 1:2000 | | |
| anti-mouse IgG-HRP | | SANTA CRUZ BIOTECHNOLOGY | 1:2000 | | |
| **d (CyTOF)** | | | | | |
| **Target Protein** | **Element** | **Isotope** | **Clone** | **Vendor** | **Concentration (ug/ml)** |
| CD45 | In | 115 | 30-F11 | Biolegend | 3 |
| pSTAT3 | La | 139 | 4 | BD | 4 |
| IgD | Ce | 140 | 11-26c.2a | BD | 2 |
| B220 | Pr | 141 | RA3-6B2 | BD | 0.5 |
| pHH3 | Nd | 142 | HTA28 | Biolegend | 4 |
| pMAPKAPK2 | Nd | 144 | 27B7 | CST | 2 |
| pCREB | Nd | 145 | 87G3 | CST | 3 |
| pPLCg2 | Nd | 146 | K86-689.37 | BD | 0.25 |
| pSTAT1 | Nd | 150 | 4a | BD | 4 |
| pErk | Eu | 151 | D13.14.4E | CST | 6 |
| Cyclin B1 | Eu | 153 | GNS-1 | Fluidigm | 1ul |
| CD27 | Sm | 154 | LG.3A10 | Biolegend | 0.5 |
| CD3 | Gd | 157 | 17A2 | BD | 2 |
| pAKT | Gd | 158 | J1-223.371 | BD | 0.375 |
| pSTAT5 | Tb | 159 | 47 | BD | 4 |
| p4EBP1 | Dy | 162 | 236B4 | CST | 2 |
| IkB | Dy | 163 | L35A5 | CST | 3 |
| pRb | Dy | 164 | J112-906 | BD | 1.5 |
| Ki67 | Ho | 165 | SolA15 | eBioscience | 3 |
| pSrc | Er | 167 | K98-37 | BD | 3 |
| p-p38 | Er | 168 | 36/p38 | BD | 3 |
| pNFkB | Tm | 169 | K10-895.12.50 | BD | 4 |
| pSyk/ZAP70 | Er | 170 | 17a | BD | 3 |
| cleaved PARP | Yb | 171 | F21-852 | BD | 4 |
| pS6 | Yb | 172 | N7-548 | Fluidigm | 1ul |
| CD19 | Yb | 173 | 6D5 | Biolegend | 1.5 |
| IgM | Yb | 174 | RMM-1 | Biolegend | 2.25 |
| CD23 | Lu | 175 | B3B4 | BD | 1.5 |
| MHC II | Yb | 176 | M5/114.15.2 | Biolegend | 0.1875 |

| Supplementary Table 2 | | |
|---|---|---|
| **Reagents** | **Cat#** | **Vendor** |
| Cisplatin | P4394 | Millipore-Sigma |
| Paraformaldehyede | 15710 | Electron Microscopy Science |
| Click-iT™ EdU Alexa Fluor™ 647 Flow Cytometry Assay Kit | C10419 | Thermo Fisher Scientific |
| Click-iT™ EdU Alexa Fluor™ 488 Flow Cytometry Assay Kit | C10425 | Thermo Fisher Scientific |
| IdU | I7125 | Millipore-Sigma |
| DAPI | D1306 | Thermo Fisher Scientific |
| Phosphoflow Perm Buffer III | 558050 | BD Biosciences |
| Perm/wash Buffer | 554723 | BD Biosciences |
| 4-(3,5-Bis(benzyloxy)phenyl)-6-(5-chloro-2-hydroxyphenyl)-2-oxo-1,2-dihydropyridine-3-carbonitrile, LLP3 | SML0991 | Millipore-Sigma |
| Palbociclib (PD0332991) Isethionate, CDK 4/6 inhibitor | S1579 | Selleck Chemicals |
| Cdk1/2 Inhibitor III - CAS 443798-55-8 CDK 2/1 inhibitor | 217714 | Millipore-Sigma |
| RPMI 1640 Medium | 11875093 | Thermo Fisher Scientific |
| Fetal Bovine Serum (FBS) heat inactivated | 100-106 | Gemini Bio-products |
| Penicillin-Streptomycin-Glutamine (100X) | 10378016 | Thermo Fisher Scientific |
| QIAzol Lysis Reagent | 79306 | Qiagen |
| miRNeasy Mini Kit | 217004 | Qiagen |
| RNase-Free DNase Set | 79254 | Qiagen |
